# Molecular and functional properties of PFC astrocytes during neuroinflammation-induced anhedonia

**DOI:** 10.1101/2020.12.27.424483

**Authors:** Blanca Diaz-Castro, Alexander M. Bernstein, Giovanni Coppola, Michael V. Sofroniew, Baljit S. Khakh

**Author notes:** Post publication corresponding authors. ***Author contributions*** BD-C carried out most of the experiments and AMB performed the behavioural work. GC provided guidance on the analysis of RNA-seq data. MVS and BSK conceived, designed and directed the project, and guided data analyses. BD-C analysed data and assembled the figures with feedback from BSK. BSK and BD-C wrote the paper and all authors commented and edited it. ***Data availability statement*** Excel file 1 contains the annotated RNA-seq data and all the raw data have been submitted to the Gene Expression Omnibus (GEO) with accession ID GSE135110. Any other data are available upon reasonable request.

## Abstract

Astrocytes are widely implicated in CNS diseases, but their contributions to disease related phenotypes remain incompletely explored. Anhedonia accompanies several neurological and psychiatric diseases, including major depressive disorder (MDD) and Alzheimer’s disease (AD), both of which are associated with neuroinflammation. In order to explore how neuroinflammation affects astrocytes, we assessed medial prefrontal cortex (PFC) and visual cortex (VCX) astrocytic gene expression using a neuroinflammation mouse model that displayed anhedonia as a phenotype. In this model, anhedonia was reversed by the fast acting antidepressant ketamine. Astrocyte specific gene expression alterations included those related to immune cell signaling, intracellular Ca^2+^ signaling, cholesterol biosynthesis, and metabolic pathways. Such changes peaked when anhedonia was greatest, and reversed to normal when anhedonia subsided. However, region-specific molecular identities between PFC and VCX astrocytes were maintained throughout, implying that astrocyte identities do not converge during neuroinflammation. We also mapped anhedonia-related astrocyte and bulk tissue gene expression changes onto published PFC single cell RNA sequencing data, and compared them to MDD and AD post-mortem human tissue samples to identify shared mechanisms. Finally, we assessed how neuroinflammation affected mPFC neuronal properties and detected no alterations at a time point when there was strong astrocyte reactivity. Our data show that neuroinflammation can cause significant and reversible changes in astrocyte gene expression and mouse behaviour without obvious neurotoxicity or loss of essential homeostatic functions. Furthermore, gene expression signatures accompanying neuroinflammation reveal pathways shared with MDD and AD, which display neuroinflammation as a comorbidity in humans.

**Significance statement:** Astrocytes are widely implicated in brain diseases, but their contributions to disease-related phenotypes remain incompletely explored. To make inroads into this problem, we assessed medial prefrontal cortex (PFC) and visual cortex (VCX) astrocyte gene expression using a peripherally induced neuroinflammation mouse model that produced anhedonia – a phenotype associated with several brain disorders. Neuroinflammation caused reversible changes in mouse behaviour and astrocyte-specific gene expression changes, some of which were related to human post mortem data for major depressive disorder (MDD) and Alzheimer’s disease (AD), but without any clear evidence of neurotoxicity in PFC of mice. The astrocyte molecular alterations accompanying neuroinflammation-induced anhedonia will be informative to explore diverse brain disorders and the effects of neuroinflammation on the CNS more broadly.

## Introduction

Neuroinflammation is a complex multicellular response that accompanies a variety of disorders including neurological and psychiatric diseases (Giovannoni and Quintana, 2020; Linnerbauer et al., 2020). In many disorders involving neuroinflammation, it has proven problematic to separate correlation from causation and thus the extent to which neuroinflammation accompanies brain diseases or is directly causal for disease-related phenotypes remains unclear. In these regards, a hypothesis based on evolutionary theories and clinical observations postulates that inflammatory dysregulation can lead to depressive behavior in humans, which may contribute to major depressive disorder (MDD) (Miller and Raison, 2016; Syed et al., 2018). This supports genetic, neuropathological, and cellular studies that link immune cell dysfunction to several neurological and psychiatric disorders (Prinz and Priller, 2014; Barnes et al., 2017; Ferrer, 2017; Lai et al., 2017; Hickman et al., 2018; Perez-Nievas and Serrano-Pozo, 2018; Pimenova et al., 2018; Pouget, 2018; Syed et al., 2018; Hammond et al., 2019; Matthews, 2019). Furthermore, there are known associations between inflammation and inflammatory markers with MDD-related phenotypes (Syed et al., 2018; van Eeden et al., 2020). Based on such discoveries, there is a need to understand how neuroinflammation affects relevant cell types in brain nuclei known to contribute to brain disorders and therefore begin to explore the significance of neuroinflammation to disease related phenotypes including, but not limited to, MDD.

Astrocytes are among the main cell types that respond to, and contribute, neuroinflammatory cues in the CNS (Sofroniew, 2014; Bennett and Molofsky, 2019). Astrocytes are the most numerous type of glia and tile the entire CNS. They intimately contact most, if not all, brain cells and the vasculature. Astrocytes participate in the formation and function of the blood-brain barrier, provide metabolic support to neurons, are involved in synapse formation and removal, and regulate neuronal activity by buffering ions and neurotransmitters, as well as through regulated mechanisms that include the release of neuroactive substances (Barres, 2008). In addition, astrocytes are altered in all forms of brain disease, injury, and trauma, and are therefore proposed to contribute in a correlative or causative manner to disease states (Burda and Sofroniew, 2014; Giovannoni and Quintana, 2020; Linnerbauer et al., 2020). Much has been learnt about basic mechanisms, but exploration of how astrocytes contribute to psychiatric disorders is still in its infancy. In this study, we sought to understand astrocyte alterations accompanying anhedonia induced by neuroinflammation in mice. We focused on anhedonia as a behavioral readout because this phenotype is strongly associated with MDD and dementia-related disorders such as Alzheimer’s disease (AD), and it is readily observed and quantified in mice (Scheggi et al., 2018; De Fruyt et al., 2020).

We performed a series of astrocyte functional evaluations, immunohistochemistry (IHC), behavioral assessments, and RNA sequencing (RNA-seq) in the mouse prefrontal cortex (PFC) of mice injected with a single peripheral dose of lipopolysaccharide (LPS) to mimic antigen presentation and neuroinflammation (Zamanian et al., 2012; Bhattacharya et al., 2018). As the main integrator of the decision making circuits of the limbic system, the PFC becomes dysfunctional in both psychiatric and dementia-related disorders and when symptoms such as anhedonia are observed (Russo and Nestler, 2013; Elahi and Miller, 2017; Dafsari and Jessen, 2020; Haber et al., 2020). Astrocyte specific and whole tissue RNA-seq was performed during the development and recovery of anhedonia with the goal of identifying molecular changes within astrocytes in an unbiased way (Srinivasan et al., 2016; Chai et al., 2017; Yu et al., 2018; Yu et al., 2020). In addition, we compared astrocyte gene expression data from PFC to that from visual cortex (VCX) to evaluate how neuroinflammation affected astrocyte regional transcriptional identity. We then mapped neuroinflammation and anhedonia-related astrocyte and bulk tissue gene expression changes onto PFC single cell RNA sequencing data, and compared them to MDD and AD post-mortem human tissue samples to identify shared mechanisms. Finally, we performed a range of physiological evaluations to determine how astrocyte reactivity that accompanied neuroinflammation affected the properties of astrocytes and neurons in the PFC.

## Materials and Methods

Most animal experiments were conducted in accordance with the National Institute of Health Guide for the Care and Use of Laboratory Animals and were approved by the Chancellor’s Animal Research Committee at the University of California, Los Angeles. A subset of the animal experiments conformed to national and institutional guidelines including the Animals [Scientific Procedures Act] 1986 (UK), and the Council Directive 2010/63EU of the European Parliament and the Council of 22 September 2010 on the protection of animals used for scientific purposes, and had full Home Office ethical approval. All mice were housed with food and water available *ad libitum* in a 12 hour light/dark environment. All animals were sacrificed during the light cycle, and none were involved in previous studies.

### Mice

Experiments were performed using 8-10 week old C57BL/6J (JAX# 000664) mice. For astrocyte specific RNA-seq experiments, B6N.129-Rpl22^tm1.1Psam^/J (JAX# 011029) (Sanz et al., 2009) were bred with *Aldh1l1*-cre/ERT2 mice (Srinivasan et al., 2016) from an in-house colony; male and female homozygous knock-in hemizygous transgenic mice were used. For Ca^2+^ imaging Ai95 (JAX# 024105) were bred with *Aldh1l1*-cre/ERT2 mice. All experiments except for the RNA-seq were performed with female mice. All mice were young adults between 7 and 9 weeks old at the time of LPS injection.

### Drug administration

Lipopolysaccharide (Sigma #L5024) from *E. coli* was diluted to 1 mg/ml in sterile PBS and stored in aliquots at −80°C. Eight to nine week old mice received one single injection of 5 mg/kg at 11 AM. Mice were weighed every day from the day of injection until sacrifice. Fluoxetine HCl (Tocris #0927) was dissolved in 100% ethanol (20 mg/ml), then mixed with sterile PBS to a concentration of 2 mg/ml and stored at −80°C. Each animal received either 10 mg/kg fluoxetine or 100 μl of sterile PBS intraperitoneally each morning for 2 weeks prior to LPS injection, continuing until day of sacrifice. Ketamine HCl (Ketalar® NDC-42023-115-10) 100 mg/ml was diluted to 2 mg/ml. Mice were injected with 10 mg/kg ketamine or sterile PBS one time intraperitoneally immediately following LPS injection. Tamoxifen (Sigma, 20 mg/ml) was administered intraperitoneally for five consecutive days at 100 mg/kg body weight to 6-7 week old *Aldh1l1*-cre/ERT2 × RiboTag mice (Rpl22^tm1.1Psam^). Mice were use 2-3 weeks after the last tamoxifen injection.

### Behavioral assessments

#### Sucrose Preference Test

One week before LPS injections, mice underwent a 3 day baseline test for sucrose preference. Each cage received two bottles filled with water *ad libitum*. The following day, one bottle was replaced by 1% sucrose solution. On day 3, that bottle was replaced by water. Consumption of each liquid was measured. One day prior to LPS injection, each cage received a fresh bottle of water and 1% sucrose solution, which were weighed every day until sacrifice. Sucrose preference was calculated by the following formula: 100*weight of the liquid drunk from the sucrose bottle/weight of the liquid drunk from the two bottles.

#### Open Field Test (OFT)

Mice injected with PBS, LPS (5 mg/kg) or those that were uninjected (as control for effects of the injection on locomotor activity), were placed in plastic containers measuring 30.5 × 18 × 25.5 cm inside a dimly lit room. Mice underwent OFT 6 hours after the injection and on days one, two, three, ten, and 20 at 5 PM. Mice were allowed to acclimate for 3 min, and then were recorded for the following 6 min. To eliminate ambient noise, a white noise generator was used (San Diego Instruments). AnyMaze video analysis software was used to quantify time immobile, time spent in the center of the arena and number of rearing bouts.

### Transcardial perfusions

For transcardial perfusion, mice were euthanized with pentobarbitol (i.p.) and perfused with 10% buffered formalin (Fisher #SF100-20). Briefly, once all reflexes subsided, the abdominal cavity was opened. The animal was perfused with 6 ml of 0.1 M phosphate buffered saline (PBS) containing 10 units/ml of heparin, followed by 60 ml 10% buffered formalin. After gentle removal from the skull, the brain was post fixed in 10% buffered formalin overnight at 4°C. The tissue was cryoprotected in 30% sucrose PBS solution the following day for at least 48 hours at 4°C until use.

### Immunohistochemistry (IHC)

Coronal sections (40 μm) of formalin perfused brains were prepared using a cryostat microtome (Leica) and kept in 0.05 M PBS, 250 mM sucrose, 7 mM MgCl_2_, 50% Glycerol at −20°C. For IHC, sections were washed twice in 0.1 M PBS for 10 min, once in 1M HCl, and then incubated in blocking solution containing 10% normal donkey or goat serum (NDS or NGS) in 0.1 M PBS with 0.05% Triton® X-100 for 1 hr at room temperature with agitation. Sections were subsequently incubated in primary antibodies diluted in 0.1 M PBS with 0.05% Triton® X-100 overnight at 4°C with agitation. The following primary antibodies were used: rat anti-GFAP (1:1000; Thermofisher 13-0300), rabbit anti-GFAP (1:1000; Dako Z0334), rabbit anti-S100ß (1:3000; Abcam Ab41548), mouse anti-HA (1:1000; Biolegend 901514), rabbit anti-NeuN (1:2000; Cell Signalling D3S3I), guinea pig anti-NeuN (1:1000, Synaptic Systems 266-004), guinea pig anti-Iba1 (1:800, Synaptic Systems 234-004). The next day the sections were washed three times in 0.1 M PBS for 10 min each before incubation at room temperature for 2 hr with secondary antibodies diluted in blocking solution. The following secondary antibodies were used: goat anti-mouse 488 (1:1000, Invitrogen A10680), goat anti-rabbit 546 (1:1000, Invitrogen A11010), donkey anti-rat 488 (1:500, Invitrogen A-21208), donkey anti-rabbit 546 (1:500, Invitrogen A10040), donkey anti-guinea pig 594 (1:500, Jackson 706-585-148), donkey anti-guinea pig 647 (1:500, Jackson 706-605-148), donkey anti-rat 647 (1:500, Jackson 712-605-150), donkey anti-rabbit 647 (1:500, Jackson 711-605-152), streptavidin conjugated Alexa 488 (1:250, Invitrogen S32354). The sections were rinsed two times in 0.1 M PBS for 10 min each, and once in 0.1 M PBS with 0.2 % Triton® X-100 and 140 ng/μl Dapi for 10 min. The sections were mounted on microscope slides in fluoromount-G. Fluorescent images were taken using either UplanSApo 20X 0.85 NA oil immersion objective lens on a commercial confocal laser-scanning microscope (Fluoview V1000; Olympus), a 25X).8 NA oil immersion lens on a Zeiss LSM900 confocal laser-scanning microscope, or a Zeiss Axioplan 2 with FluoArc pseudo-confocal. The settings were the same within each experiment.

### RNA-seq of mouse whole tissue and astrocyte transcriptomes

Mice were anesthetized with isoflurane and decapitated. The brain was quickly extracted and cut into 2 mm coronal slices using a rodent brain matrix (Electron Microscopy Sciences). Slices were placed on ice cold PBS and PFC and VCX were further dissected and collected for RNA extraction. Whole tissue or astrocyte specific RNA was used for RNA-seq studies. For whole tissue PFC, C57/BL6J mice were used. The dissected tissue was homogenized and RNA was purified following the RNeasy kit instructions (RNeasy Plus Micro QIAGEN #74034). For determining the astrocyte transcriptomes Ribotag mice crossed with *Aldh1l1*-cre/ERT2 mice were used. Briefly, freshly dissected tissues were collected and individually homogenized in 1 ml of homogenization buffer (50 mM Tris pH 7.4, 100 mM KCl, 12 mM MgCl2, 1% NP-40, 1 mM Dithiothreitol (DTT), 1X Protease inhibitors, 200 U/ml RNAsin, 100 μg/ml Cyclohexamide, 1 mg/ml Heparin). RNA was extracted from 20% of cleared lysate as input (200 μl). The remaining lysate (800 μl) was incubated with 5 μl of mouse anti-HA antibody (Biolegend #901514) with rocking for 4 hours at 4°C followed by addition of 200 μl of magnetic beads (Pierce #88803) and overnight incubation with rocking at 4°C. The beads were washed three times in high salt solution (50 mM Tris pH 7.4, 300 mM KCl, 12 mM MgCl_2_, 1% NP-40, 1 mM Dithiothreitol (DTT), 100 μg/ml Cyclohexamide). RNA was purified from the IP and corresponding input samples (RNeasy Plus Micro QIAGEN #74034).

RNA concentration and quality were assessed with an Agilent 2100 Bioanalyzer. RNA samples with RNA integrity numbers (RIN) greater than seven (mean RIN 8.5) were processed with Ribo-Zero Gold kit (Epicentre, WI) to remove ribosomal RNA. Sequencing libraries were prepared using Illumina TruSeq RNA sample prep kits following the manufacturer’s protocol. After library preparation, amplified double-stranded cDNA was fragmented into 125 bp (Covaris-S2, Woburn, MA) DNA fragments, which were (200 ng) end-repaired to generate blunt ends with 5’-phosphates and 3’-hydroxyls and adapters ligated. The purified cDNA library products were evaluated using the Agilent Bioanalyzer (Santa Rosa, CA) and diluted to 10 nM for cluster generation in situ on the HiSeq paired-end flow cell using the CBot automated cluster generation system. Four separate sequencing experiments were performed, one for whole tissue PFC 1 d after injection, another one for IP and INPUT 1 d PFC and VCX, a third one for IP and INPUT 2 d PFC and VCX, and a fourth one for IP and INPUT 14 d PFC and VCX. Samples in each experiment were multiplexed into a single pool in order to avoid batch effects and 69 bp-paired-end sequencing was performed using an Illumina HiSeq 4000 sequencer (Illumina, San Diego, CA). A yield between 30 and 90 million reads was obtained per sample.

Quality control was performed on base qualities and nucleotide composition of sequences. Alignment to the *M. musculus* (mm10) refSeq (refFlat) reference gene annotation was performed using the STAR spliced read aligner with default parameters. Additional QC was performed after the alignment to examine: the degree of mismatch rate, mapping rate to the whole genome, repeats, chromosomes, key transcriptomic regions (exons, introns, UTRs, genes), insert sizes, AT/GC dropout, transcript coverage and GC bias. Total counts of read-fragments aligned to candidate gene regions were derived using HTSeq program (www.huber.embl.de/users/anders/HTSeq/doc/overview.html) with mouse mm10 refSeq (refFlat table) as a reference and used as a basis for the quantification of gene expression. Only uniquely mapped reads were used for subsequent analyses. Four mice per condition were used; however, in the 2 d dataset two outlier samples were excluded, one LPS-PFC-IP, and another one LPS-VCX-INPUT. Differential expression analysis was conducted with R-project and the Bioconductor package limma-voom. RNA-seq data has been deposited within the Gene Expression Omnibus (GEO) repository (www.ncbi.nlm.nih.gov/geo), with accession number GSE135110.

### Astrocyte intracellular Lucifer yellow iontophoresis

This method for filling cells in fixed tissue was modified from published methods (Bushong et al., 2002; Chai et al., 2017). C57BL/6J mice injected with PBS or LPS were euthanized two days after the injection with pentobarbitol (i.p.) overdose and transcardially perfused with 10 ml of 35°C Ringer’s Solution with 0.02% lidocaine followed by 50 ml of 10% buffered formalin (Fisher #SF100-20). Ringer’s Solution contains the following (in mM): 135 NaCl, 14.7 NaHCO_3_, 5 KCl, 1.25 Na_2_HPO_4_, 2 CaCl_2_, 1 MgCl_2_, and 11 D-glucose, bubbled with 95% O_2_ and 5% CO_2_. Brains were lightly post-fixed at room temperature for 1.5 hr and then washed three times in ice cold 0.1 M PBS for 10 min. 100 μm coronal sections were cut using Pelco Vibrotome 3000 and then placed in ice-cold PBS for the duration of the experiment. 10 mg Lucifer yellow CH di-Lithium salt (Sigma) was dissolved in 1 ml 5 mM KCl solution and filtered prior to use. Sharp (200 MOhm) glass electrodes were pulled from Borosilicate glass capillarie with filament (O.D. 1.0 mm, I.D. 0.58 mm). Electrodes were gravity filled. Sections were transferred to a solution of room temperature PBS for filling. Astrocytes were identified using IR-DIC and then impaled with the sharp electrode. Lucifer yellow was injected into the cell by applying ~ 1 volt for 10 minutes. The electrode was removed and the filled astrocyte was imaged using a confocal microscope (Fluoview V1000; Olympus), with UplanFL 40X water-immersion objective lens 0.8 NA and at a digital zoom of two to three and a 0.25 um confocal z-steps. Quantitative soma and branch + process volumes were generated using Imaris’ surface function. The astrocyte territory volume was measured from a low-intensity threshold reconstruction encompassing the cell volume and the space between its processes. The number of primary branches was counted visually.

### Acute brain slice preparation for imaging and electrophysiology

Slice procedures have been described previously (Shigetomi et al., 2013; Jiang et al., 2016). Coronal striatal or hippocampal slices were prepared from 8-9 week old mice two days after injection of PBS or LPS. Briefly, animals were deeply anesthetized with isoflurane and decapitated. The brains were placed and sliced in ice-cold modified artificial cerebrospinal fluid (aCSF) containing the following (in mM): 194 sucrose, 30 NaCl, 4.5 KCl, 1 MgCl_2_, 26 NaHCO_3_, 1.2 NaH_2_PO_4_, and 10 D-glucose, saturated with 95% O_2_ and 5% CO_2_. A vibratome (DSK-Zero1) was used to cut 300 μm brain sections. The slices were allowed to equilibrate for 30 min at 32-34°C in normal aCSF containing (in mM); 124 NaCl, 4.5 KCl, 2 CaCl_2_, 1 MgCl_2_, 26 NaHCO_3_, 1.2 NaH_2_PO_4_, and 10 D-glucose continuously bubbled with 95% O_2_ and 5% CO_2_. Slices were then stored at 21–23°C in the same buffer. All experiments were performed within 4-6 hr of slicing.

### Electrophysiological recording and assessment of cell coupling in brain slices

Slices were placed in the recording chamber and continuously perfused with 95% O_2_ and 5% CO_2_ bubbled normal aCSF. Cells were visualized with infrared optics on an upright microscope (Fluoview FV1000; Olympus). pCLAMP10 software and a Multi-Clamp 700B amplifier was used for electrophysiology (Molecular Devices). For recording from astrocytes and dye coupling experiments, currents were measured in whole-cell mode using pipettes with a typical resistance of 5.5 MΩ when filled with internal solution containing the following (in mM): 130 K-gluconate, 2 MgCl_2_, 10 HEPES, 5 EGTA, 2 Na-ATP, 0.5 CaCl_2_, with pH set to 7.3. Cells were recorded in normal aCSF and in aCSF + 250 μM Ba^2+^. For a subset of cells, 2 mg/ml biocytin was added to the intracellular solution to examine gap junction coupling. Astrocytes were held in whole-cell mode for 20 min to allow biocytin to diffuse from the patched cell to other cells connected by gap junctions. In some cases carbenoxolone (CBX; 100 μM) was added to the recording solution to block gap junctions. Brain slices were then rescued from the recording chamber for IHC.

For recording from pyramidal neurons, currents were measured in whole-cell mode using pipettes with a typical resistance of 5–6 MΩ when filled a K^+^ internal solution consisting of the following (in mM): 135 potassium gluconate, 5 KCl, 0.5 CaCl_2_, 5 HEPES, 5 EGTA, 2 Mg-ATP and 0.3 Na-GTP, pH 7.3 adjusted with KOH. In some cases, 2 mg/ml biocytin was added to the intracellular solution to subsequently visualize the patched neuron. Neurons were voltage-clamped at −70 mV unless otherwise stated. Excitatory postsynaptic potentials (EPSCs) were recorded in the presence of bicuculline (10 mM), and mini-EPSCs in bicuculline (10 μM) and tetrodoxin (TTX) (300 nM). ClampFit 10.7 software was used to analyze traces from astrocyte and neuronal recordings.

### Intracellular Ca^2+^ imaging

Slice preparation was performed as above. Cells for all the experiments were imaged using a confocal microscope (Fluoview FV1000; Olympus) with a 40X water-immersion objective lens with a numerical aperture (NA) of 0.8. A 488 nm line of an Argon laser was used, with the intensity adjusted to 9% of the maximum output of 10 mW and 2-3 digital zoom. The emitted light pathway consisted of an emission high pass filter (505-525 nm) before the photomultiplier tube. Astrocyte somata were typically 25 μm below the slice surface and scanned at 1 frame per second for imaging sessions. Spontaneous signals were recorded in aCSF + 300 nM TTX. For evoked Ca^2+^ signals, slices were washed on 100 μM adenosine 5’ triphosphate (ATP) or 10 μM phenylephrine (PE) for 1 min.

### Experimental design and statistical analyses

#### RNA-seq differential expression analyses

Statistical significance of the differential expression was determined at false discovery rate (FDR) < 0.05. In the cases in which an FPKM > 1 cutoff was used, we excluded the genes that had an FPKM < 1 in at least one of the samples being compared. Gene ontology analyses were performed using Ingenuity Pathway Analysis (IPA – QIAGEN Bioinformatics) with *P*-value < 0.05 and PANTHER overrepresentation test with FDR < 0.05.

#### INPUT combined with sc-RNAseq analysis

Mouse frontal cortex sc-RNAseq gene expression data was obtained from http://dropviz.org/ (Saunders et al., 2018). We obtained a list of cell enriched genes per cell type identified in this dataset by comparing each cell type to the rest and selecting a “minimum fold ratio” of 2, “maximum P-value exponent” of −2, “min mean log amount in target” of 0.25 and a “max mean log amount in comp” of 2. We compared the list of cell enriched genes to our list of DEGs in INPUT 1 d and 2 d NIM vs control and assigned some of the DEGs to the cells in which they were enriched. To find cell-cell interaction molecules, we first identified the IPA pathways that were common between cell types. Within those, we looked for DEGs that encode for receptors, ligands or adhesion molecules. We further filtered these by keeping the ones that were enriched in less than two cells types based on our analysis of Saunders et.al. 2018. With this information, we identified the potential cell-cell interactions mechanisms that could be altered. For example, in the IPA “Neuroinflammation signaling pathway”, we found the receptor Csf1r is upregulated and enriched in microglia. We checked if its potential ligand were cell enriched, and found that its ligand Csf1, was enriched and also upregulated in astrocytes. We used IPA to identify cell enriched upstream regulators. To detect potential non-cell autonomous regulation, we compared the list of upstream regulators and selected the ones that were cell specific, based on our analysis of Saunders et al., 2018.

#### Comparison of NIM RNA-seq with human MDD and AD sn-RNAseq

A list of sn-RNA-seq cell type specific DEGs from human PFC of MD (Nagy et al., 2020) and AD (Mathys et al., 2019) patients was obtained from their respective manuscript publications. The DEGs that had a mouse ortholog were selected and compared to the DEGs of our mouse 1 d and 2 d PFC NIM vs control analysis. For astrocyte genes, the comparison was made with the IP DEGs; for microglia, with the INPUT DEGs. PANTHER gene ontology analysis was used to identify the biological processes that were represented by genes that changed in both human disease and mouse NIM.

#### Immunohistochemistry analysis

All images were analyzed with Fiji (NIH). Cell counting was performed from maximum intensity projections using the Cell Counter plugin; only cells with somata completely within the region of interest (ROI) were counted. For signal area and intensity measurements, images were converted to 8-bit and then standardized ROIs were created and thresholded to eliminate background noise in experimental and control images. Two ROIs were analyzed per section (1 per hemisphere), and 3 sections were analyzed per animal in both PFC and VCX.

#### Calcium imaging analysis

Analyses of time-lapse image series were performed using Fiji (NIH). XY drift was corrected using a custom plugin for Fiji; cells with Z-drift were excluded from analyses. The data were analyzed essentially as previously reported (Yu et al., 2018). Time traces of fluorescence intensity were extracted from the soma, branch or process ROI signals in Fiji and presented as the change in fluorescence relative to the baseline (dF/F). A signal was declared as a Ca^2+^ transient if it exceeded the baseline by greater than twice the baseline noise. Peak amplitude, half-width, frequency, time of the first peak and area under the curve of Ca^2+^ signals were analyzed in ClampFit10.7 (Molecular Devices).

#### General data analyses and statistics

Statistical tests were run in GraphPad Instat 3. Summary data are presented as mean ± s.e.m. Note that in some of the graphs, the bars representing the s.e.m. are smaller than the symbols used to represent the mean. For each set of data to be compared, we determined within GraphPad Instat whether the data were normally distributed or not. If they were normally distributed, we used parametric tests. If the data were not normally distributed, we used non-parametric tests. Paired and unpaired Student’s two-tailed *t* tests (as appropriate) and two-tailed Mann–Whitney tests were used for most statistical analyses with significance declared at *P <* 0.05, but stated in each case with a precise *P* value. When the *P* value was less than 0.0001, it is stated as such to save space on the figure panels and text. *N* is defined as the numbers of cells or mice throughout on a case-by-case basis depending on the particular experiment; the unit of analysis is stated in each figure or figure legend. A statistical FDR value < 0.05 was used for all RNA-seq analyses. Where appropriate, key statistics are also reported in the text. No data points were excluded from any experiment.

## Results

### Neuroinflammation induced anhedonia was ketamine-sensitive

To study the effects of inflammation on the CNS, we adapted a peripherally induced neuroinflammation model, which we abbreviate as NIM for NeuroInflammation Model (Rodrigues et al., 2018). We injected a single dose of lipopolysaccharide (LPS; 5 mg/kg) intraperitoneally (i.p.) without direct perturbation of the CNS (Bhattacharya et al., 2018). For all the experiments reported herein, control mice were handled equivalently and received an injection of PBS (Fig. 1A). Subsequently, general health status and a focused set of behaviors were assessed over 20 days. The mice were weighed and anhedonia was evaluated daily with the sucrose preference test (Scheggi et al., 2018). Locomotor activity was assessed using open field tests at 1, 2, 3, 10, and 20 days after LPS injection (Fig. 1A-D, Extended data Fig. 1A-E). We found that weight loss in NIM mice peaked between days 2 and 3, and then slowly recovered back to normal by day 10 (Extended data Fig. 1B, n = 8 mice). Sucrose preference was also significantly reduced in the NIM mice for up to two days after LPS, returning back to control levels on day 3 (Fig. 1B, Extended data Fig. 1D,E; n = 8 mice). In contrast, locomotor activity was only significantly reduced on day 1 (Extended data Fig. 1C; n = 8).

**Figure 1.**
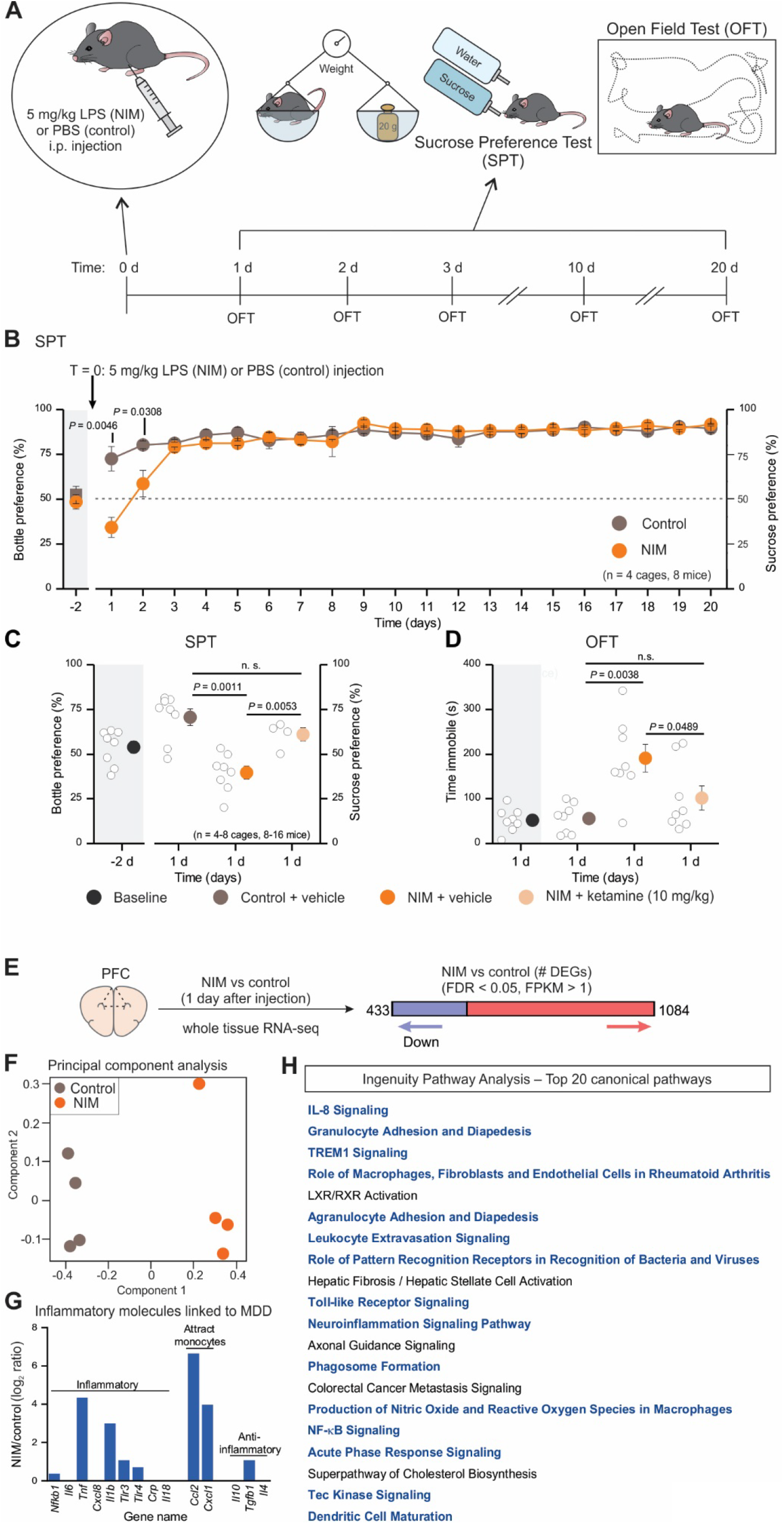
LPS induced anhedonia, sickness behavior, and overall inflammatory responses. ***A,*** Timeline for the neuroinflammation model (NIM) and its behavioral assessments. ***B,*** Sucrose preference test (SPT) results during 20 days after LPS or PBS i.p. injections. ****C,**** SPT one day after PBS, LPS or LPS + ketamine injection. Grey shading on the left of the graphs indicates the bottle preference when the mice were given the choice between two bottles of water. ****D,**** Time immobile during 6 minutes of open field test (OFT). ****E,**** One day after LPS or PBS i.p. injection, whole tissue PFC RNA-seq detected 1517 differentially expressed genes (DEGs) between NIM and control, with FDR < 0.05 and FPKM > 1. ****F,**** Principal component analysis of the top 2000 most variable genes. ****G,**** NIM vs control log_2_ ratio of the main inflammatory molecules that previously have been associated with MDD (Miller and Raison, 2016). ****H,**** Top 20 pathways altered in NIM PFC astrocytes, identified by Ingenuity Pathway Analysis. The pathways involved in inflammatory response are highlighted in blue. Grey shading on the left of each graph, indicates baseline conditions from non-injected mice. Open circles are raw data with closed circles indicating mean ± s.e.m. In some cases, the error bars representing s.e.m are smaller than the symbol used for the mean.

Since anhedonia is observed in MDD and in mouse models of MDD, we determined which of the behavioral assessments reported in the previous section were altered by two drugs used to treat MDD. Subanesthetic doses of ketamine induce rapid and long-lasting antidepressant responses in MDD and in mouse models (Berman et al., 2000; Walker et al., 2013). Notably, a single i.p. injection of 10 mg/kg ketamine (after NIM induction) rescued the anhedonia phenotype and locomotor activity in the open field (Fig. 1C,D, Extended data Fig. 2; n = 8 mice). In contrast to the effects of ketamine, the selective serotonin reuptake inhibitor (SSRI) fluoxetine injected at 10 mg/kg every day (i.p.) for two weeks before the LPS injection and subsequently for the duration of the experiment, did not significantly affect any of the LPS-evoked changes in behavior (Extended data Fig. 2). To the extent that ketamine acts as a rapid antidepressant and that it also rescued LPS-evoked changes in anhedonia, we subsequently interpreted anhedonia in mice as an MDD-related behavioral phenotype and sought to assess the accompanying astrocyte molecular changes. Ketamine-insensitive responses following LPS likely represent general sickness and were beyond the scope of this study.

Next, to evaluate if peripheral LPS did indeed cause inflammatory responses in the PFC, we performed bulk RNA-seq for four NIM and four control mice. 1517 Differentially expressed genes (DEGs) were found between NIM and control PFC one day after LPS injections (Fig. 1E). Principal component analysis (PCA) separation of the two treatments by component 1 corroborated clear transcriptional differences between NIM and control samples (Fig. 1F). Within the DEGs, we found upregulation of several inflammatory molecules (Fig. 1G) associated with MDD (Miller and Raison, 2016) and Ingenuity Pathway Analysis (IPA) showed that 15 of the top 20 altered pathways in the PFC were related to inflammatory responses (Fig. 1H, in blue). In addition, there were other interesting altered pathways such as cholesterol biosynthesis (Fig. 1H). Thus, peripheral administration of LPS induced anhedonia, which was rescued by ketamine, and was associated with clear neuroinflammatory transcriptional profiles in the PFC (Fig. 1).

### RiboTag to assess mPFC and VCX astrocyte-specific transcriptional changes

To evaluate PFC astrocyte gene expression changes during neuroinflammation, we performed astrocyte specific RNA-seq by crossing RiboTag mice (Sanz et al., 2009) to astrocyte specific *Aldh1l1*-cre/ERT2 mice (Srinivasan et al., 2016; Chai et al., 2017). Bigenic progeny were injected with tamoxifen at six weeks of age to induce the expression of Rpl22HA (RiboTag) specifically in astrocytes (Extended data Fig. 3A-E). Three weeks later, the mice were injected with LPS (or PBS) to trigger neuroinflammation. Whole tissue (INPUT) and astrocyte specific RNA (IP) was extracted for PFC as well as VCX and sequenced (Extended data Fig. 3F). We performed such analyses 1, 2 and 14 days after induction of neuroinflammation. The astrocyte-specific IP fraction was replete with known astrocyte markers (e.g. *Aldh1l1*, *Aldoc*, *Slca2*, *Gja1*) and depleted of markers of other cells for PFC and VCX datasets at all three time points (Extended data Fig. 3G,H). Furthermore, 100 markers known to be enriched in cortical astrocytes (Srinivasan et al., 2016) were also enriched in the PFC and VCX IP fractions, but depleted from the input samples (Extended data Fig. 3I). Similarly, 100 transcripts known to be depleted from cortical astrocytes were enriched in the input fractions, and depleted in the IP fractions (Extended data Fig. 3J). Thus, we could reliably assess astrocyte specific gene expression 1, 2 and 14 days after the induction of neuroinflammation.

### Impact of neuroinflammation on PFC and VCX astrocyte transcriptional identity

To address how transcriptional changes related to the behavioral alterations reported in Fig. 1, we analyzed RNA-seq of INPUT and IP samples at three time points in the NIM and control mice. We focused on Day 1 when both locomotor dysfunction and anhedonia were observed, Day 2 when locomotor activity was normal, but anhedonia persisted, and Day 14 when there were no significant differences between the NIM and control groups in the measures we assessed (Fig. 1, Extended data Fig. 1). PFC and VCX gene expression was analyzed to determine if differences existed between these two functionally distinct cortical areas. Initial assessment of the transcriptional differences between IP samples showed that the NIM group was separated from controls by the first principal component at Days 1 and 2, but not at Day 14 (Fig. 2A-C). PCA also revealed clear transcriptional differences between astrocytes from PFC and VCX, implying diversity between these areas under control conditions that persisted following neuroinflammation at Day 1, 2 and 14 (Fig. 2A-C). Accordingly, we found 1218 differentially expressed genes (DEGs) between PFC and VCX regions in control conditions (Fig. 2D), and 1176 DEGs in NIM conditions (Fig. 2E). To evaluate the genes more directly, we compared the PFC enriched genes (yellow in Fig. 2D,E) from control and NIM samples. We found that more than half of the PFC enriched genes did not change their region specificity following neuroinflammation (Fig. 2F), and thus appeared intrinsic to these astrocytes irrespective of the neuroinflammatory insult. A similar trend was observed for the VCX enriched genes (green in Fig. 2D,E), implying that neuroinflammation does not markedly cause astrocytes to lose their PFC and VCX regional molecular identities (Fig. 2G). The top 20 PFC and VCX enriched genes are listed in Fig. 2H,I, respectively. Some top PFC enriched genes were transcription factors (*Zic3* and *Zic1*) or involved in cell growth and proliferation (*Greb1*, *Acvr1c*, *Zic1*, *Nrn1*). We had previously identified μ-crystallin a protein enriched in striatal astrocytes (Chai et al., 2017). Interestingly, another crystallin (*Crybb1*) was identified as enriched in PFC vs VCX astrocytes. *Crybb1* has been linked to anxiety and schizophrenia (Spadaro et al., 2015). Several of the VCX enriched genes were related to the formation and function of the extracellular matrix (*Colq*, *Fgfbp1*, *Spp1*, *Egfl6*). Cell adhesion molecules were found in both PFC and VCX astrocytes: *Bves* was enriched in PFC, *Cdh1* in VCX, for example.

**Figure 2.**
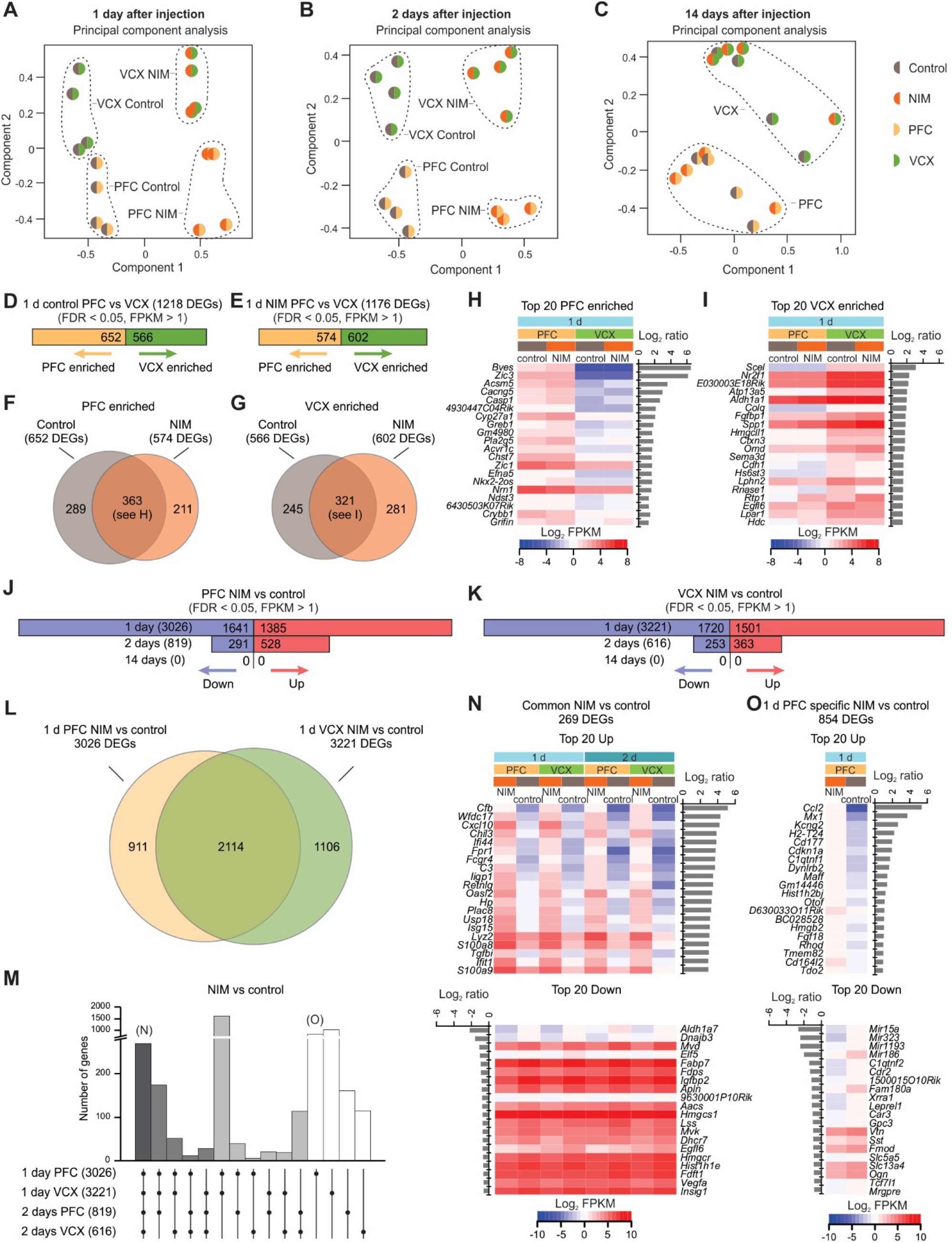
Cortical region-specific and neuroinflammation time-specific transcriptional alterations in astrocytes. ***A-C**,* Principal component analysis (top 1000 most variable genes) of NIM and control astrocytes RNA, from PFC and VCX, one (*A*), two (*B*) and 14 days (*C*) after LPS or PBS injection. *D,* Number of astrocyte DEGs between 1 d control PFC and VCX (FDR < 0.05, FPKM > 1). *E,* Number of astrocyte DEGs between 1 d NIM PFC and VCX (FDR < 0.05, FPKM > 1). *F,* Overlap of PFC enriched genes (compared to VCX) between 1d control and NIM conditions. *G,* Overlap of VCX enriched genes (compared to PFC) between 1d control and NIM conditions. *H,* Top 20 PFC enriched genes when compared to VCX in both 1 d control and NIM conditions (see *F*). *I,* Top 20 VCX enriched genes when compared to PFC in both 1 d control and NIM conditions (see *G*). *J-K,* Number of PFC (*J*) or VCX (*K*) NIM vs control DEGs (FDR < 0.05, FPKM > 1) in astrocytes at 1 d, 2 d and 14 d. *L,* Overlap of 1 d NIM vs control DEGs between PFC and VCX (FDR < 0.05, FPKM > 1). *M,* Overlap of NIM vs control astrocyte DEGs (FDR < 0.05, FPKM > 1) between PFC and VCX at 1 d and 2 d. *N,* Top 20 NIM vs control upregulated (top) and downregulated (bottom) genes that are common across 1 d PFC, 1 d VCX, 2 d PFC and 2 d VCX astrocytes. *O,* Top 20 NIM vs control upregulated (top) and downregulated (bottom) genes that are uniquely altered (FDR < 0.05, FPKM > 1), in 1 d PFC astrocytes.

We next focused on PFC and VCX astrocyte gene expression changes in response to neuroinflammation (Fig. 2J,K). Both PFC and VCX astrocytes showed large transcriptional responses at Day 1, with > 3000 DEGs. This number was reduced to < 1000 DEGs at Day 2, and remarkably no significant gene expression changes were observed at Day 14 (with an FDR < 0.05 threshold). Since we observed PFC and VCX astrocytes were transcriptionally separable, we tested if their responses to neuroinflammation were also different. Focusing on Day 1, we found that 2114 genes of the 3026 and 3221 DEGs were altered in both PFC and VCX, respectively (Fig. 2L). Of these 2114, 269 DEGs maintained their differences at Day 2 (Fig. 2M,N). Upregulated genes in this highly conserved group of genes are mostly involved in inflammatory/immune responses and the downregulated genes were related to metabolism (Fig. 2N, Extended data Fig. 4B). Notably, up to one third of the significantly DEGs caused by neuroinflammation were specific to the brain region (Fig. 2L,M). Broadly, these data demonstrate that astrocytes from different brain areas show diverse responses to the same stimulus and underscore the necessity of drawing conclusions based on evaluations of defined brain areas, rather than mixing between brain regions. These data add support to the emerging view that astrocytes are separable between brain areas and respond in a context-specific manner (Chai et al., 2017; Haim and Rowitch, 2017; Khakh and Deneen, 2019; Yu et al., 2020).

In the Day 1 dataset, 854 genes were unique to the PFC (Fig. 2M): of these, the top 20 up and downregulated DEGs are shown in Fig. 2O. Evaluation of these genes shows that region and time specific DEGs were involved in a wide variety of functions (Extended data Fig. 4). Within the pathways altered at Day 1 in PFC astrocytes of particular interest were those related to “positive regulation of neuron projection development”, “cell morphogenesis involved in neuron differentiation” and the inhibition of “actin cytoskeleton signaling” (Extended data Fig. 4C). Moreover, specific analyses of known functions performed by astrocytes suggested potential alterations of K^+^ homeostasis (Extended data Fig. 5A), Ca^2+^ signaling (Extended data Fig. 5B), gap junctions (Extended data Fig. 5C), as well as synapse morphogenesis and removal (Extended data Fig. 5D). Taken together the aforementioned RNA-seq data reveal astrocyte diversity that was dependent on the brain area and the time from induction of neuroinflammation. In particular, the data show quantitatively that NIM presented profound changes in gene expression at Day 1, and that these were completely reversible by Day 14. Furthermore, even though neuroinflammation evoked strong changes in gene expression, astrocytes maintained many PFC *versus* VCX regional gene expression differences.

### Exploring brain cell type contributions to neuroinflammation

The totality of the mechanisms by which a peripheral stimulus such as LPS results in changed behavior are unknown, but likely involve multiple cell types. We next used the INPUT RNA-seq data, which represents all cells, to explore this aspect. Recalling observations with astrocyte specific data (Fig. 2), the first two principal components of the PCA plot separated control from NIM mice, and PFC from VCX at Day 1 and Day 2. At Day 14, there was no separation between NIM and control, but there was for PFC and VCX as observed in controls (Fig. 3A-C). Almost 2000 DEGs between NIM and control mice were detected at Day 1 for PFC and VCX, but this value fell to ~400 DEGs at Day 2, and to zero at Day 14 (Fig. 3D,E). In order to assign these DEGs to their likely cell types, we took advantage of recently published single cell (sc) RNA-seq data that identified 14 cell types in the mouse PFC (Saunders et al., 2018) (Fig. 3F). Using that dataset (http://dropviz.org/) we generated a list of enriched genes for each of the 14 cell types, and determined which were differentially expressed between NIM and control mice (Fig. 3G,H, Extended data Fig. 6). Based on the overlap with the scRNA-seq data, 515 of the 1868 DEGs at Day 1 could be mapped to specific cell types (Fig. 3G). Most downregulated genes mapped to neurons, followed by astrocytes as the second largest cell type. In contrast, the upregulated genes mapped to microglia and endothelial cells. For the Day 2 time point, 178 of the 412 PFC DEGs were assigned to specific cell types (Fig. 3H). Only 13 were downregulated and most mapped to astrocytes. Two thirds of the upregulated genes at Day 2 mapped to microglia, followed by endothelial cells. To evaluate how many of the DEGs at Day 2 were shared with those at Day 1, we calculated the overlap between them based on cell-type (Fig. 3I,J). Thus, 62-64% of the DEGs observed at Day 2 for microglia, endothelial cells and astrocytes were already significantly differentially expressed on Day 1 (neurons and oligodendrocytes displayed only two DEGs on Day 2). Interestingly, all cell types showed a strong decrease in the number of DEGs from Day 1 to 2, with the exception of microglia (Fig. 3J,K). By mapping our INPUT RNA-seq data onto cell types known to exist in the PFC, our analysis demonstrates that neuroinflammation involves a multicellular response that wanes from Day 1 to Day 2 in most cells (including astrocytes) except microglia.

**Figure 3.**
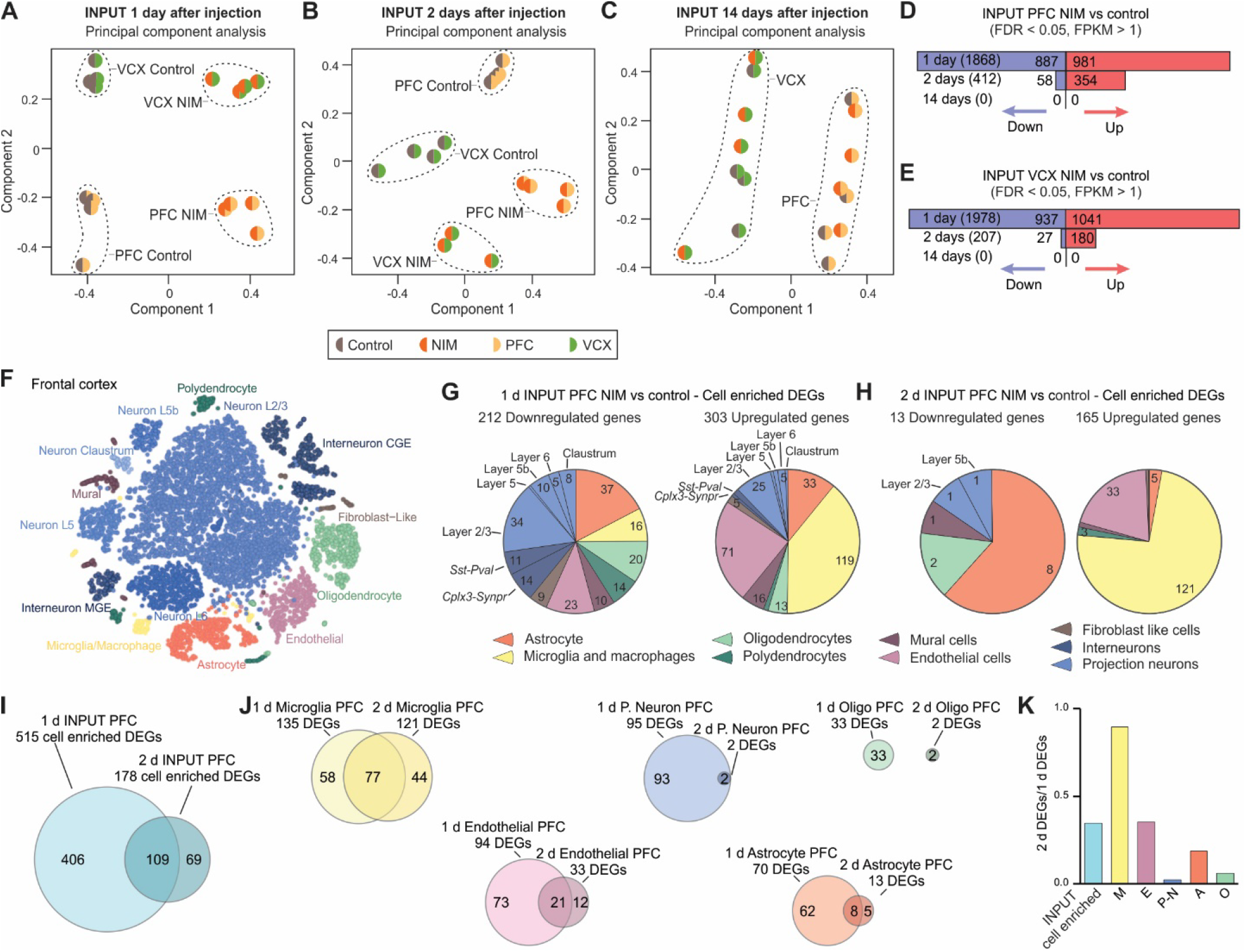
Assessment of PFC cell specific responses during neuroinflammation from input RNA-seq. ***A-C,*** Principal component analysis (top 2000 most variable genes) of NIM and control input RNA, from PFC and VCX, one (*A*), two (*B*) and 14 days (*C*) after LPS/PBS injection. *D-F,* Number of PFC (*D*) or VCX (*E*) NIM vs control DEGs (FDR < 0.05, FPKM > 1) in input samples at 1 d, 2 d and 14 d. *F,* tSNE plot of the mouse frontal cortex main cell types identified with single cell (sc)RNA-seq (http://dropviz.org/) (Saunders et al., 2018). *G,* Pie charts indicating the proportion of 1 d input downregulated (left) and upregulated (right) genes that are enriched in each cell type listed in the legend below. The numbers in each section of the pie chart indicate the number of DEGs for that cell type. *H,* Same as in g but for 2 d input samples. *I,* Overlap of all cell enriched NIM vs control DEGs between 1 d and 2 d. *J,* Overlap of microglia, endothelial, projection neuron (p-neuron), astrocyte and oligodendrocyte NIM vs control DEGs between 1 d and 2 d. *K,* Bar graph representing the 2 d/1 d ratio of the number of DEGs for all cell enriched genes (input), microglia (M), endothelial (E), projection neuron (P-N), astrocyte (A) and oligodendrocyte (O).

### Brain cell type responses and cell-cell interactions during neuroinflammation

We used IPA to explore the pathways that were altered in microglia, endothelial cells, neurons, astrocytes, and oligodendrocytes (Extended data Fig. 7A-O). Microglial pathways indicated direct engagement of these cells in immune responses and phagocytosis (Extended data Fig. 7A,B). Endothelial cells were involved in inflammatory and vascular signaling (Extended data Fig. 7D,E). Projection-neuron altered pathways related to stress responses, analgesia and neuronal function (Extended data Fig. 7G,H). Pathways affected in astrocytes were mostly metabolic (Extended data Fig. 7J,K). Oligodendrocytes underwent metabolic and other diverse changes (Extended data Fig. 7M,N). Comparison of the pathways identified at Day 1 and Day 2 for each cell type showed that even though the DEGs at each stage vary (Fig. 3J), microglia and endothelial cells maintain their functional roles at these time points (Extended data Fig. 7C,F), whereas other cells do so to a lesser degree (Extended data Fig. 7I,L,O).

**Table 1:** Summary and statistics for astrocyte intracellular calcium imaging data.

| Astrocyte compartment | Parameter | Sample | Average | Median | SEM | n | P value | Statistical test |
| --- | --- | --- | --- | --- | --- | --- | --- | --- |
| Soma | Peak amplitude (dF/F) | Control | 1.3 | 0.4 | 0.2 | 57 events, 15 cells, 4 mice | 0.0438 | Mann Whitney |
|  |  | NIM | 1.0 | 0.2 | 0.2 | 65 events, 20 cells, 5 mice |  |  |
|  | Half-width (s) | Control | 5.0 | 3.8 | 0.5 | 55 events, 15 cells, 4 mice | 0.0742 | Mann Whitney |
|  |  | NIM | 4.2 | 2.9 | 0.4 | 61 events, 20 cells, 5 mice |  |  |
|  | Frequency (events/min) | Control | 0.6 | 0.5 | 0.1 | 20 cells, 4 mice | 0.4857 | Mann Whitney |
|  |  | NIM | 0.5 | 0.4 | 0.1 | 29 cells, 5 mice |  |  |
| Branches | Peak amplitude (dF/F) | Control | 0.7 | 0.5 | 0.1 | 137 events, 19 cells, 4 mice | 0.0002 | Mann Whitney |
|  |  | NIM | 0.6 | 0.3 | 0.04 | 199 events, 28 cells, 5 mice |  |  |
|  | Half-width (s) | Control | 3.6 | 3.0 | 0.3 | 121 events, 19 cells, 4 mice | 0.0390 | Mann Whitney |
|  |  | NIM | 3.0 | 2.3 | 0.2 | 173 events, 28 cells, 5 mice |  |  |
|  | Frequency (events/min) | Control | 1.4 | 1.0 | 0.4 | 19 cells, 4 mice | 0.9408 | Mann Whitney |
|  |  | NIM | 1.4 | 0.8 | 0.2 | 29 cells, 5 mice |  |  |
| Territory | Peak amplitude (dF/F) | Control | 0.1 | 0.1 | 0.004 | 190 events, 18 cells, 4 mice | 0.4244 | Mann Whitney |
|  |  | NIM | 0.1 | 0.1 | 0.003 | 400 events, 28 cells, 5 mice |  |  |
|  | Half-width (s) | Control | 6.5 | 5.1 | 0.3 | 149 events, 18 cells, 4 mice | 0.1563 | Mann Whitney |
|  |  | NIM | 6.0 | 4.9 | 0.2 | 293 events, 28 cells, 5 mice |  |  |
|  | Frequency (events/min) | Control | 1.9 | 2.2 | 0.2 | 20 cells, 4 mice | 0.0209 | Unpaired T-test |
|  |  | NIM | 2.8 | 2.8 | 0.3 | 29 cells, 5 mice |  |  |
| Treatment | Parameter | Sample | Average | Median | SEM | n | P value | Statistical test |
| ATP (100 μM) | Time of the first peak (s) | Control | 84 | 81 | 4 | 16 cells, 4 mice | 0.9059 | Unpaired t test with Welch correction |
|  |  | NIM | 85 | 84 | 2 | 14 cells, 4 mice |  |  |
|  | Area under the curve (dF/F*s) | Control | 130 | 104 | 16 | 16 cells, 4 mice | 0.9534 | Mann Whitney |
|  |  | NIM | 134 | 138 | 14 | 15 cells, 4 mice |  |  |
| PE (10 μM) | Time of the first peak (s) | Control | 71 | 66 | 5 | 17 cells, 4 mice | 0.8363 | Mann Whitney |
|  |  | NIM | 67 | 67 | 1 | 17 cells, 4 mice |  |  |
|  | Area under the curve (dF/F*s) | Control | 282 | 278 | 20 | 17 cells, 4 mice | 0.8355 | Unpaired t test with Welch correction |
|  |  | NIM | 286 | 294 | 8 | 17 cells, 4 mice |  |  |

Note to table: the time of the first peak for the ATP and PE-evoked responses is the time taken for the response to occur after applying the agonists in the bath at 2 ml/min.

We observed that many pathways overlapped between cell types (Extended data Fig. 8), which may suggest synergistic or reciprocal cell-cell interaction mechanisms, in which, for instance, a receptor involved in a specific pathway is expressed in one cell type, and its ligand in another. We explored this possibility by focusing on common pathways between cell types and sought known cell-cell interactions that were enriched in specific cell types. We thus identified potential homotypic cell-cell interaction mechanisms that were altered during neuroinflammation (Extended data Fig. 7P). For example, several cell adhesion molecules were upregulated in NIM when compared to controls: *Cldn5* and *Pecam1* in endothelial cells or *Cdh19* in astrocytes and oligodendrocytes. Integrin pathways, with different combinations of ligands and receptors depending on the cell type, were affected in microglia and astrocytes. Interestingly, both *Sst* within interneurons and its receptors *Sstr2* and *Sstr3* in projection neurons were downregulated. Somatostatin interneuron deficiency has been proposed as a mechanism involved in depression (Fee et al., 2017) and somatostatin infusion induces antidepressant like effects (Engin and Treit, 2009). Heterotypic interactions were also found to be changed (Extended data Fig. 7Q). We identified several upregulated pathways whereby astrocytes, endothelial cells and oligodendrocytes produced ligands that signaled to microglia. For example, *Csf1r* in microglia was found to be upregulated in NIM, its ligand *Csf1*, was enriched in astrocytes and oligodendrocytes (Saunders et al., 2018). *Csf1r* has been identified to be essential for microglia survival (Elmore et al., 2014). Integrin heterotypic communication was also detected between microglia and astrocytes or endothelial cells. We next used IPA to identify predicted upstream regulators that may influence the expression of the DEGs in different cell types (Extended data Fig. 7R).

### Comparison of PFC transcriptional profiles of NIM with human MDD and AD

It has been suggested that astrocyte neuroinflammatory phenotypes have fundamental roles in both neuropsychiatric and neurodegenerative diseases (Sofroniew, 2015; Liddelow and Barres, 2017; Wang et al., 2017; Perez-Nievas and Serrano-Pozo, 2018). We compared our NIM data to recent post-mortem human PFC single nuclei (sn) RNA-seq from MDD and AD patients (Mathys et al., 2019; Nagy et al., 2020) to assess if the astrocyte neuroinflammatory responses we observed in NIM were reproduced within human disease. Nagy and colleagues compared single nuclei transcriptomes of the PFC of 17 MDD patients and 17 controls (Nagy et al., 2020). They identified 7 DEGs in astrocytes (Fig. 4A), three of them were also altered at Day 1 in PFC NIM astrocytes (Fig. 4B). From these, *Fads2*, a gene encoding for a protein involved in fatty acid biosynthesis, was downregulated in both human MDD and our NIM datasets. *Myo5a* and *Tmsb4x*, involved in actin transport and polymerization, changed in opposite directions (Fig. 4C,D).

**Figure 4.**
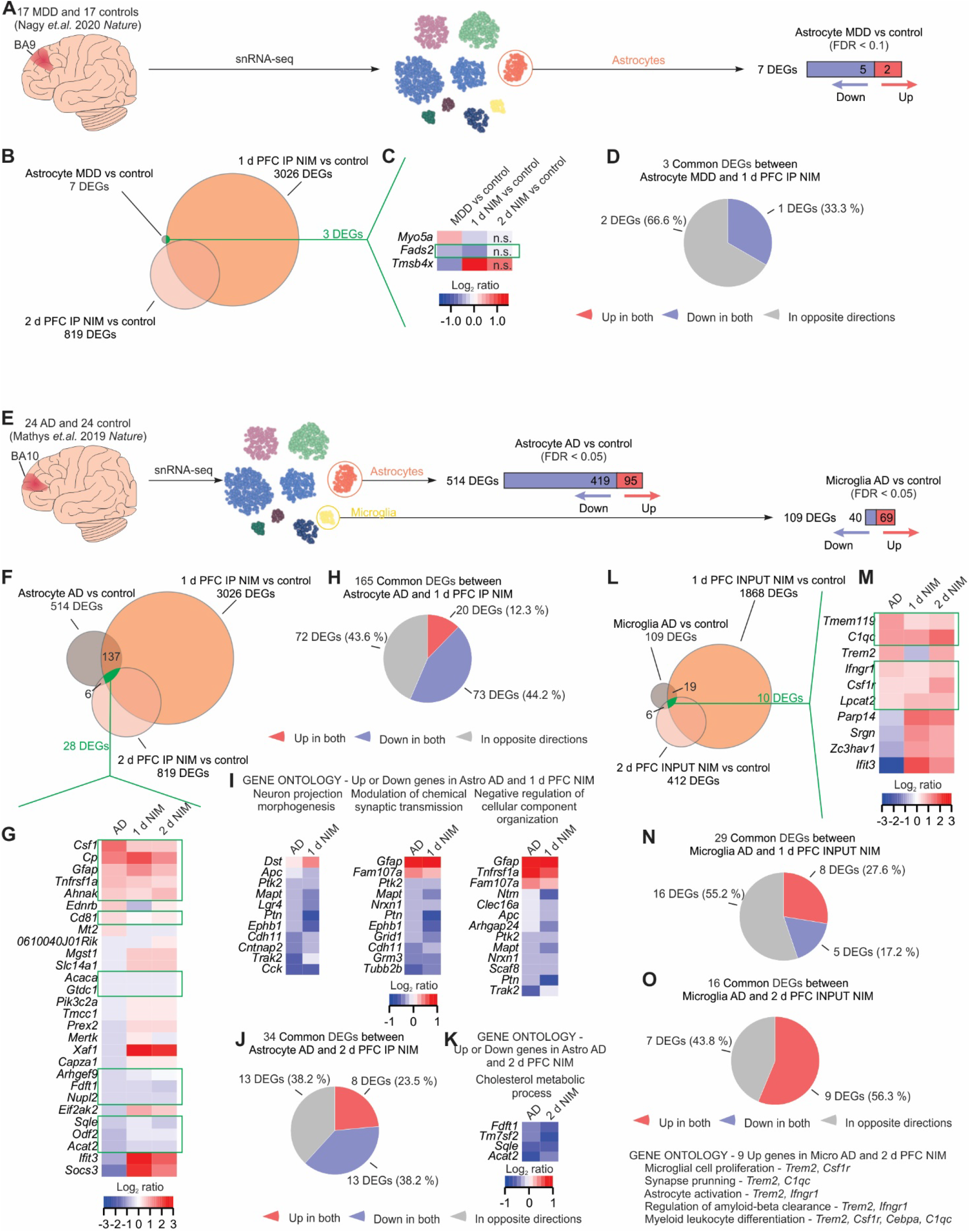
Comparison of PFC transcriptional profiles between neuroinflammation and human MDD and AD. ***A,*** Prefrontal cortex snRNA-seq data from 17 MDD patients and 17 controls identified 7 astrocyte DEGs (Nagy et al., 2020). ****B,**** Venn diagram representing the overlap between astrocyte MDD, 1 d NIM and 2 d NIM DEGs. ****C,**** Heatmap showing the differential expression magnitude in log_2_ ratio of the 3 common astrocyte DEGs between MDD and 1 d NIM. ****D,**** Pie chart indicating the agreement between astrocyte MDD and 1 d NIM on the direction of the gene expression change. ****E,**** Prefrontal cortex snRNA-seq data from 24 AD patients and 24 controls identified 514 astrocyte and 109 microglia DEGs (Mathys et al., 2019). ****F,**** Venn diagram representing the overlap between astrocyte AD, 1 d NIM and 2 d NIM DEGs. ****G,**** Heatmap showing the differential expression magnitude in log_2_ ratio of the 28 common astrocyte DEGs between AD, 1 d NIM and 2 d NIM. ****H,**** 165 genes were differentially expressed in both AD and 1 d NIM astrocytes. The pie chart indicates the agreement between astrocyte AD and 1 d NIM on the direction of the gene expression change. ****I,**** PANTHER GO analysis of the genes that changed in the same direction in both AD and 1 d NIM. The heatmaps indicate the genes included in each one of the biological process GO terms and the magnitude of their differential expression in log_2_ ratio. ****J,**** 34 genes were differentially expressed in both AD and 2 d NIM astrocytes. The pie chart indicates the agreement between astrocyte AD and 2 d NIM on the direction of the gene expression change. ****K,**** PANTHER GO analysis of the genes that changed in the same direction in both AD and 2 d NIM. The heatmaps indicate the genes included in the biological process GO term and the magnitude of their differential expression in log_2_ ratio. ****L,**** Venn diagram representing the overlap between microglia AD, 1 d NIM and 2 d NIM DEGs. ****M,**** Heatmap showing the differential expression magnitude in log_2_ ratio of the 10 common microglia DEGs between AD, 1 d NIM and 2 d NIM. ****N,**** 29 genes were differentially expressed in both AD and 1 d NIM microglia. The pie chart indicates the agreement between microglia AD and 1 d NIM on the direction of the gene expression change. ****O,**** 16 genes were differentially expressed in both AD and 2 d NIM microglia. The pie chart indicates the agreement between microglia AD and 2 d NIM on the direction of the gene expression change. Below, PANTHER GO analysis for biological process terms of the genes that changed in the same direction in both AD and 2 d NIM. In the heat maps, red means upregulation and blue downregulation when comparing the condition of study vs its control. The green frames on the heatmaps highlight the genes that change in the same direction in the human disease and mouse neuroinflammation.

Mathys and colleagues used snRNA-seq to assess PFC cell transcriptional alterations in 24 AD patients and 24 controls (Mathys et al., 2019). 524 DEGs were identified in astrocytes (Fig. 4E). From these, 28 were also altered at Day 1 and 2 in NIM PFC astrocytes (Fig. 4F,G). Furthermore, half of them changed in the same direction in both AD and NIM. The top common upregulated gene was *Csf1*, the gene that encodes the cytokine ligand of the essential microglial receptor *Csf1r* (Elmore et al., 2014), and one of the genes identified by our heterotypic cell-cell interaction analysis (Extended data Fig. 7Q). Common upregulated genes also included *Gfap* and the inflammatory mediators *Tnfrsf1a* and *Cd81*. Several of the genes downregulated in the three datasets were involved with lipid metabolism, e.g. *Acaca*, *Fdft1*, *Sqle*, and *Acat2* (Fig. 4G). 165 DEGs were altered in both AD and Day 1 NIM PFC astrocytes (Fig. 4F; Extended data Fig. 8A,B). Of these, 56% changed in the same direction in both datasets – most of them being upregulated (Fig. 4H). We performed gene ontology analyses to identify the main pathways that were altered in both AD and at Day 1 in NIM (Fig. 4I). Unexpectedly, the three biological processes identified were not related to inflammatory processes, but instead, they included neuron projection morphogenesis, modulation of synaptic transmission and cellular component organization. Several of the genes in these pathways encode cell adhesion molecules (*Cdh11*, *Cntnap2*, *Nrxn1*), receptors or ligands (*Ephb1*, *Cck*, *Ptn*, *Grid1*, *Grm3*, *Tnfrsf1a*), or cytoskeleton related proteins (*Dst*, *Gfap*, *Fam107a*, *Mapt*, *Tubb2b*) (Fig. 4I). 34 DEGs were common between AD and Day 2 NIM astrocytes (Fig. 4F; Extended data Fig. 8A,C); 61% of them changed in the same direction (Fig. 4J). Gene ontology analysis of these emphasized that cholesterol synthesis is downregulated in both AD and NIM (Fig. 4K).

We next used our INPUT data to assess how the AD microglia DEGs compared to our NIM dataset. 109 DEGs were identified in PFC of AD patients when compared to the controls (Fig. 4E). 10 of them were also altered at Day 1 and 2 in NIM PFC (Fig. 4L,M). Notably, the receptor for the astrocyte *Csf1* gene product, *Csf1r*, was one of the genes upregulated in all AD, Day 1 NIM, and Day 2 NIM microglia (Fig. 4M). Other shared genes that encode receptors included *Tmem119* and *Ifngr1*. Interestingly, *Trem2*, an AD risk gene that is upregulated in AD microglia, was downregulated at Day 1, but upregulated at Day 2 in the NIM INPUT samples. 29 DEGs were common between AD microglia and Day 1 NIM INPUT (Fig. 4N). Almost a half of them changed in the same direction. No significant gene ontology pathways were identified with this dataset. 16 DEGs were common between AD and Day 2 NIM (Fig. 4O). All pathways that changed in the same direction were upregulated and were involved in a wide variety of biological processes, e.g. proliferation, synapse pruning, astrocyte activation and amyloid-beta clearance.

To summarize, we have compared PFC transcriptional changes in response to neuroinflammation to the transcriptional changes occurring in related brain regions of two diseases that present with anhedonia (MDD and AD). Given that the number of astrocyte DEGs in the MDD dataset was very low, no major similarities were found. However, the comparison to AD were of interest. Approximately, 32 and 7% of the genes that were altered in astrocytes or microglia of AD patients were differentially expressed in the same direction in NIM at Day 1 and 2, respectively. In astrocytes, the common genes were mainly involved in immune responses, reduced support to neurons, and reduced lipid synthesis.

### Effects of neuroinflammation on astrocyte phenotypes

It has been proposed that LPS activates microglial release of cytokines that convert astrocytes into a neurotoxic A1 type which can kill surrounding neurons (Liddelow et al., 2017). We next asked if PFC and VCX astrocytes preferentially expressed the proposed pan reactive, A1 and A2 reactivity genes in NIM mice relative to controls (Zamanian et al., 2012; Liddelow et al., 2017). The majority of pan reactive and A1 genes were upregulated at Day 1 in the NIM mice for both PFC and VCX. The number of upregulated genes decreased at Day 2 and these transcriptional changes were fully recovered by Day 14 when no significant differences were observed (Fig. 5A). Upregulation of some A2 genes was also observed on Day 1 in both PFC and VCX astrocytes (Fig. 5A), which is consistent with overall astrocyte reactivity. We also performed IHC for GFAP. In accord with the gene expression data, upregulation of GFAP was observed in astrocytes of PFC and VCX (Fig. 5C-D), while the number of S100β positive astrocytes or Iba1 expressing microglia did not change (Fig. 5F-I). Although the IHC data need to be interpreted cautiously, together with RNA-seq they nonetheless suggest the existence of neuroinflammatory signatures in whole tissue (Fig. 1E-H, 5A-C; Extended data Fig. 4).

**Figure 5.**
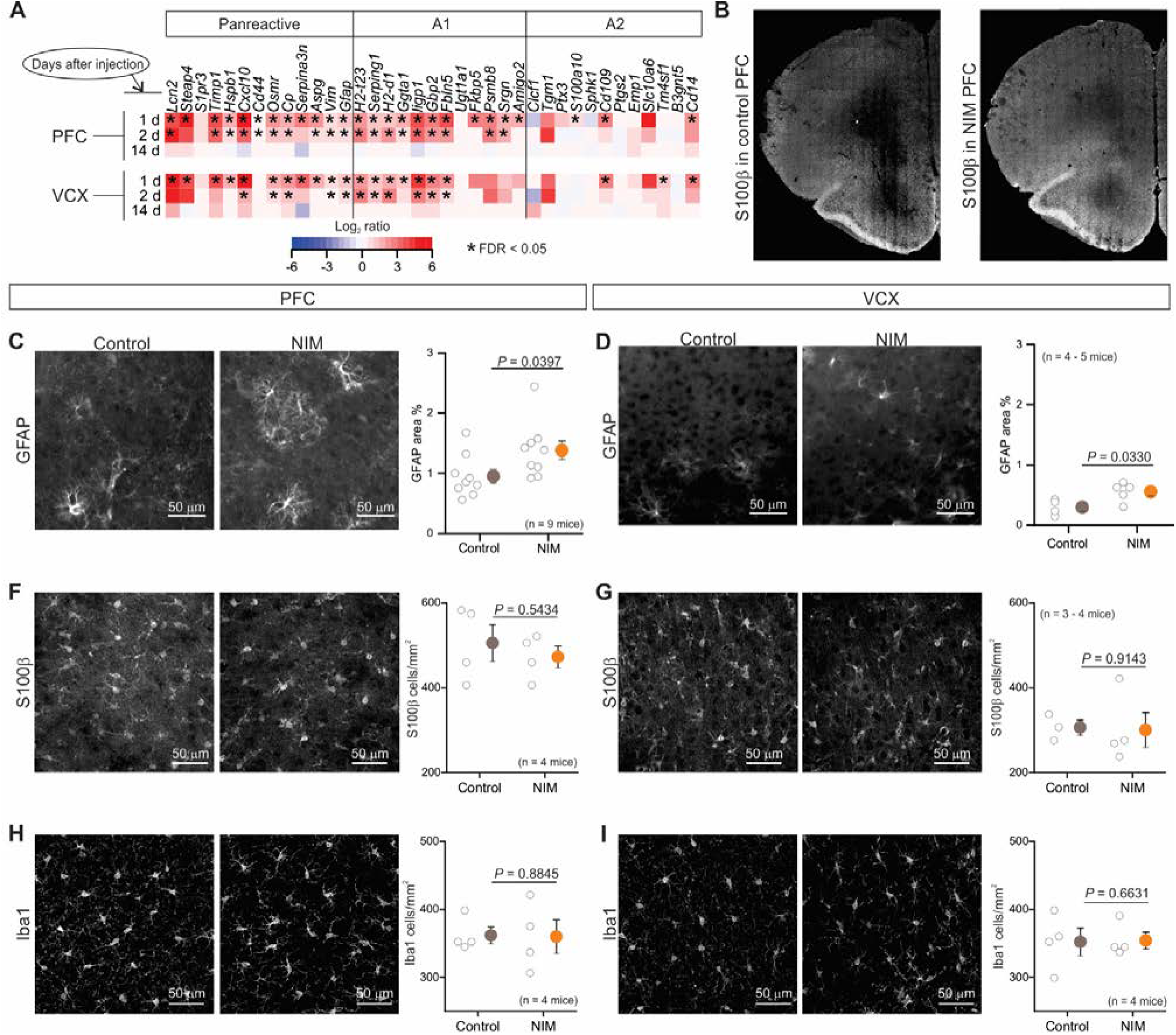
Astrocyte reactivity in PFC and VCX astrocytes. (A) NIM vs control RNA-seq differential expression of the top astrocyte reactivity genes in 1 d, 2 d and 14 d PFC and VCX astrocytes. In the heat maps, red means upregulation and blue downregulation when comparing NIM vs control. * indicates the genes that were differentially expressed with FDR < 0.05. (B) Representative immunostaining of S100β in PFC and VCX brain slices. (C,D) Representative images from GFAP staining and area % coverage of GFAP in control and NIM PFC (C) or VCX (D). (E,F) Representative images from S100β staining and number of S100β positive cells per mm^2^ in control and NIM PFC (E) or VCX (F). (G,H) Representative images from Iba1 staining and number of Iba1 positive cells per mm^2^ in control and NIM PFC (G) or VCX (H). In the scatter plots, open circles are raw data with closed circles indicating mean ± s.e.m. In some cases, the error bars representing s.e.m are smaller than the symbol used for the mean.

Since RNA-seq data identified significant changes in astrocyte transcriptional profiles, we performed a series of experiments under identical conditions to assess functional alterations that accompany reactive astrocytes in NIM versus control mice. Using intracellular Lucifer yellow iontophoresis we assessed astrocyte morphology (Fig. 6A-E), and with whole-cell patch-clamp we evaluated astrocyte resting membrane potentials, membrane conductance, current waveforms, macroscopic current-voltage relations, and Ba^2+^-sensitive currents (Fig. 6F-J). Using imaging of intracellular Ca^2+^ signals, we assessed spontaneous Ca^2+^ signaling in astrocyte somata, branches and processes, as well as GPCR-mediated Ca^2+^ signals evoked by ATP and phenylephrine, which activate P2Y and α1 adrenoceptors on astrocytes (Fig. 6K-M; Table 1). Using patch-clamp mediated intracellular loading with biocytin we monitored gap-junctional dye coupling between astrocytes (Extended data Fig. 9a). From all these assessments, we detected ~25% decrease in the astrocyte branch and process volumes following neuroinflammation (Fig. 6) and modest alterations in amplitude, half-with and frequency of spontaneous Ca^2+^ signals (Table 1). Overall, our data show that mPFC astrocytes display strong changes in gene expression when assessed agnostically, and some significant functional alterations. We emphasize that the topic of astrocyte functional changes should be revisited in the future when a greater number of metrics can be assessed reliably, because our data do not rule out the existence of significant functional changes that we could not assess carefully in our experiments. In these regards, the metrics to assess function are limited in relation to the large number of genes that can be assessed with RNA-seq.

**Figure 6.**
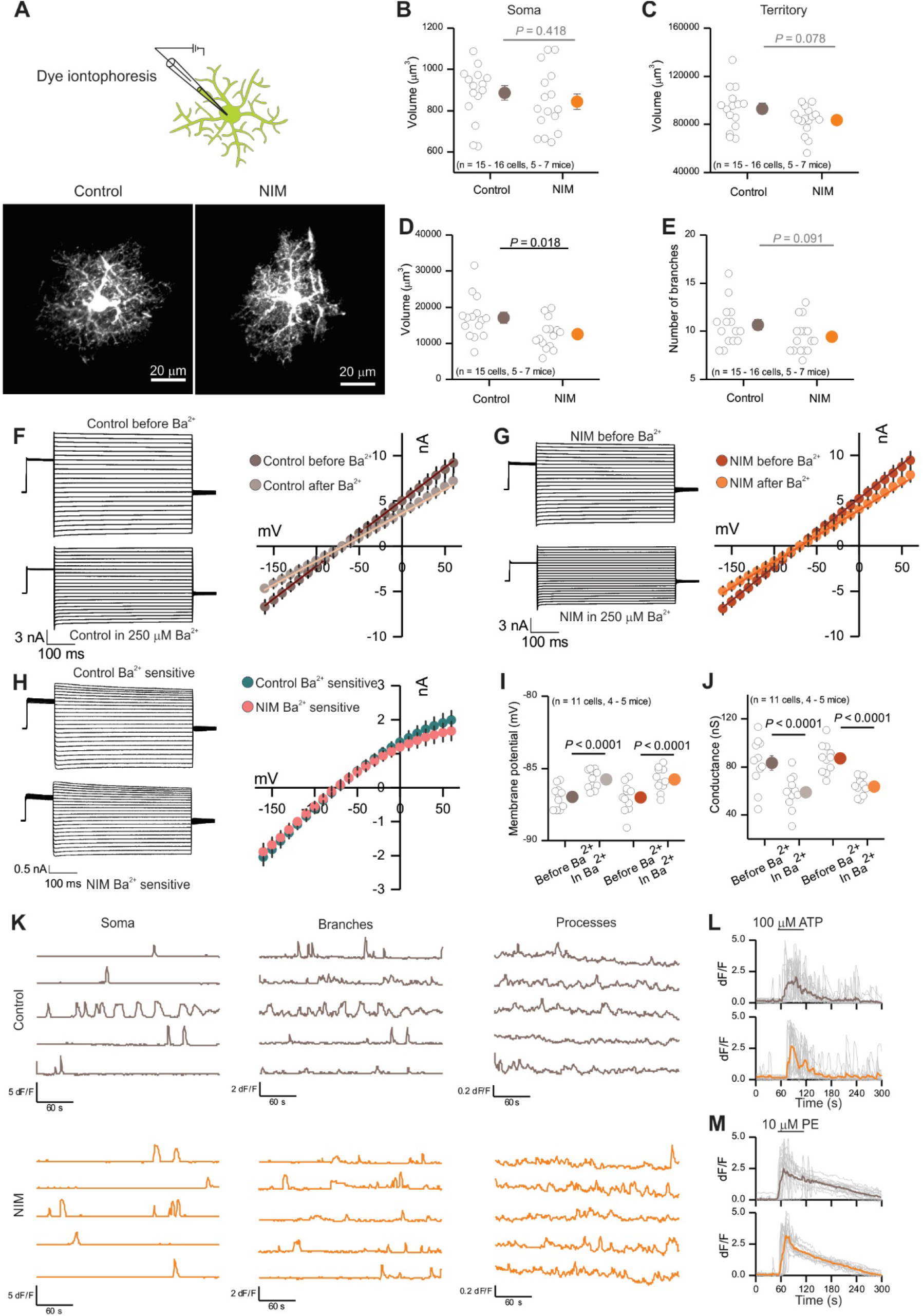
Functional and morphological alterations of PFC astrocytes during neuroinflammation. *A,* Representative images of 2 d control and NIM PFC astrocytes filled by Lucifer yellow iontophoresis. *B-D,* Volume of soma (*B*), territory (*C*) and branches + processes (*D*) of 2 d control and NIM astrocytes. Cell compartment volumes were calculated with Imaris software. ****E,**** Number of branches that ramify from the soma of 2 d control and NIM astrocytes. ****F-G,**** Whole-cell voltage clamp was performed on 2 d control (****F****) and NIM (****G****) astrocytes before and in the presence of 250 μM Ba^2+^. On the left, representative waveforms for total and Ba^2+^ insensitive currents are shown. On the right, average current-voltage relations. ****H,**** On the left, representative waveforms for control and NIM Ba^2+^ sensitive currents are shown. On the right, average current-voltage relations for control and NIM Ba^2+^ sensitive currents. ****I,**** Membrane potential of control and NIM astrocytes before and during Ba^2+^. ****J,**** Conductance of control and NIM astrocytes before and during Ba^2+^. ****K,**** Representative traces of 2 d control and NIM spontaneous Ca^2+^ signals (in 300 nM TTX) in soma, branches and processes. The amplitude of Ca^2+^ signals is bigger in control somas and branches than in NIM, however Ca^2+^ signals are more frequent in the processes of NIM than in control (Table 1). ****L-M,**** Single (grey) and average (brown or orange) traces of astrocyte soma and branch Ca^2+^ increase in response to 1 min exposure to 100 μM ATP (****L****) or 10 μM phenylephrine (PE) (****M****). In the scatter plots, open circles are raw data with closed circles indicating mean ± s.e.m. In some cases, the error bars representing s.e.m are smaller than the symbol used for the mean.

### Functional assessments of PFC neurons following neuroinflammation

To test if astrocyte reactive phenotypes in NIM had an effect on the survival of neurons, we performed IHC with the neuronal marker NeuN. No differences in neuron numbers were found between control and NIM in PFC or VCX (Fig. 7A-C). We further assessed potential alterations in neuronal excitability by performing whole-cell patch clamp of PFC pyramidal neurons of layers 2/3 (Fig. 7D-U). Remarkably, no differences between NIM and control were observed in membrane potential (Vm), membrane resistance (Rm) or the minimum current necessary to produce an action potential (rheobase) (Fig. 7D-I). Furthermore, no untoward issues were noted when recording from neurons from NIM mice and the cells looked healthy in both NIM and control conditions. We next recorded and quantified spontaneous excitatory postsynaptic currents (sEPSCs; in 10 μM bicuculline). Neither the amplitude nor the frequency of the events was different between the experimental conditions (Fig. 7J-O). Action potential independent miniature EPSCs (mEPSCs; in 10 μM bicuculline + 300 nM TTX), however, were slightly increased in frequency in the NIM neurons. There were no differences in mEPSC amplitude (Fig. 7P-U). All in all, PFC pyramidal neurons were ostensibly normal with no evidence of dysfunction or neurotoxicity under conditions when astrocyte reactivity genes associated with A1 phenotypes were significantly elevated.

**Figure 7.**
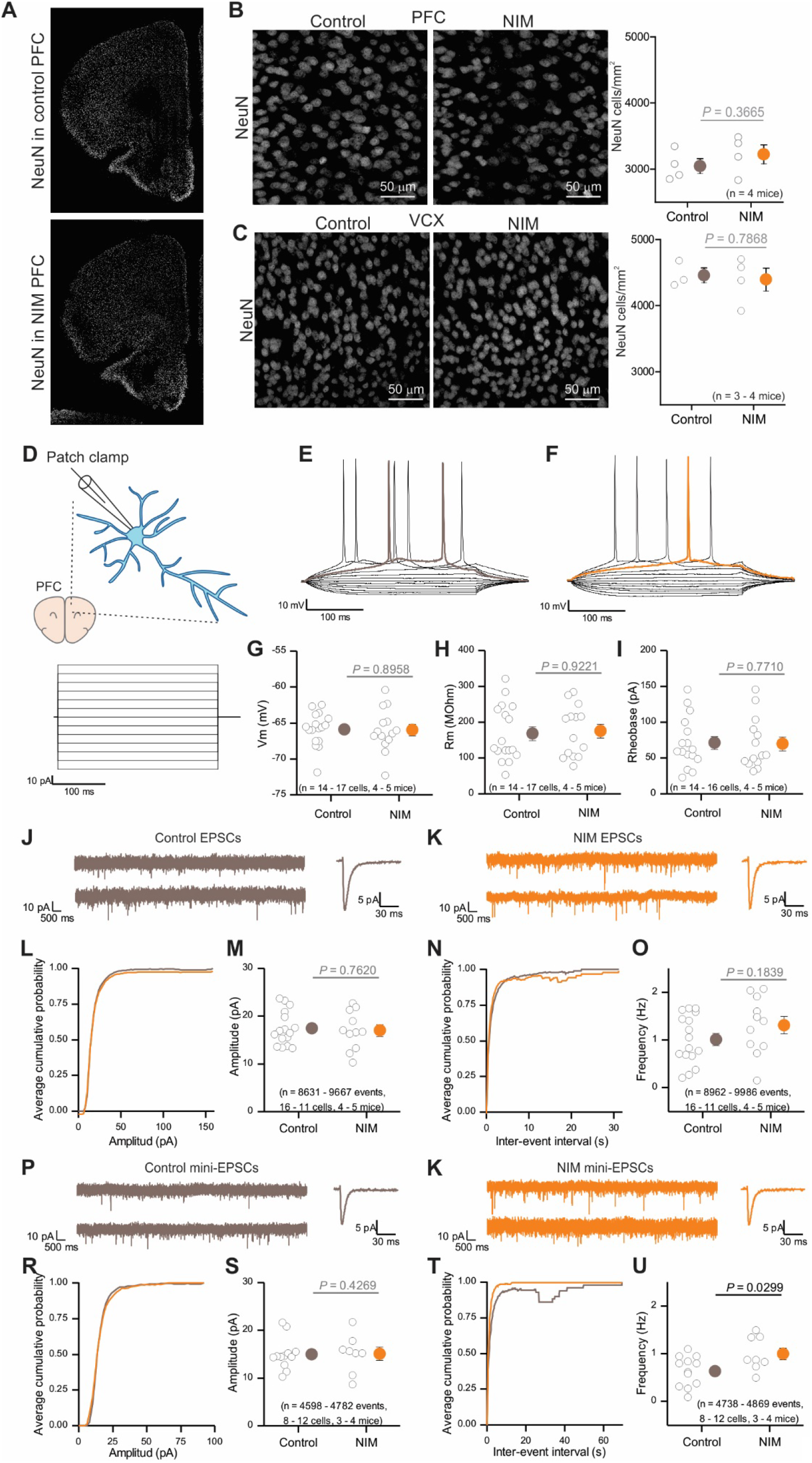
Neuronal and synaptic properties of PFC pyramidal neurons during neuroinflammation. ***A,*** Representative immunostaining of NeuN in PFC and VCX brain slices. ****B,C,**** Representative images from NeuN staining and number of NeuN positive cells per mm^2^ in control and NIM PFC (****B****) or VCX (****C****). ****D,**** Whole-cell current clamp was performed in 2 d NIM and control PFC pyramidal neurons. 10 pA steps were applied during 300 ms. ****E,F,**** Representative voltage waveforms from control (****E****) and NIM (****F****). Rheobase trace is highlighted in brown for control and orange for NIM. ****G-I,**** control and NIM pyramidal neuron membrane potential (****G****), membrane resistance (****H****), and rheobase (****I****). ****J,K,**** Representative EPSC (10 μM bicuculline) current clamp recording and single EPSC example from control (****J****) and NIM (****K****) PFC pyramidal neurons. ****L,**** Average cumulative probability plot for EPSC amplitude. ****M,**** EPSC average amplitude for 2 d control and NIM PFC pyramidal neurons. ****N,**** Average cumulative probability plot for EPSC inter-event intervals. ****O,**** EPSCs frequency for 2 d control and NIM PFC pyramidal neurons in a 10 minute recording. ****P,Q,**** Representative mini-EPSC (10 μM bicuculline and 300 nM TTX) current clamp recording and single mini-EPSC example from control (****P****) and NIM (****Q****) PFC pyramidal neurons. ****R,**** Average cumulative probability plot for mini-EPSC amplitude. ****S,**** Mini-EPSCs average amplitude for 2 d control and NIM PFC pyramidal neurons. ****T,**** Average cumulative probability plot for mini-EPSC inter-event intervals. ****U,**** Mini-EPSCs frequency for 2 d control and NIM PFC pyramidal neurons in a 10 minute recording.

## Discussion

Neuroinflammation has long been associated with CNS disorders and is emerging as a target for treatment, but many open questions remain at the interface between the immune system and the brain. In this study, we used a peripherally induced neuroinflammation model (NIM) to explore the molecular and functional profiles of astrocytes at specific behaviorally defined stages. We examined three time points: Day 1 when the mice displayed locomotor dysfunction and anhedonia, Day 2 when locomotor activity was normal, but anhedonia persisted, and Day 14 when there were no detectable differences between the NIM and controls. We performed astrocyte specific and whole tissue RNA-seq from PFC and VCX at each time point and comment on the major conclusions that can be drawn.

First, astrocyte specific RNA-seq experiments showed transcriptional diversity between PFC and VCX astrocytes in control conditions as well as during neuroinflammation at Days 1 and 2. Up to one third of DEGs between NIM and control mice were specific to the cortical region. Second, RNA-seq analyses suggested that, in addition to activation of immune signaling in astrocytes, astrocyte pathways involved in homeostatic roles such cholesterol metabolism, cytoskeleton dynamics, gap junctions, K^+^ and Ca^2+^ transport, and synapse plasticity were significantly affected in the PFC. Third, whole tissue RNA-seq analysis of the cell-type enriched markers revealed that microglia and brain endothelial cells were the main responders in NIM in both PFC and VCX. Fourth, we found that gene expression changes observed in NIM were completely reversible and no significant differences persisted at Day 14. Fifth, an increase in astrocyte transcriptome inflammatory profiles and a decrease in transcriptome profiles of some gene ontology-predicted neuron supportive functions were found in common between NIM and human AD astrocytes phenotypes. Sixth, in NIM, although astrocytes upregulated a set of genes related to astrocyte reactivity, these changes did not exert detectable detrimental effects on neuronal health or survival, indicating that astrocyte reactivity can be associated with maintenance of essential neural homeostatic functions.

Rodent models of neuroinflammation display depression-related behaviors such as anhedonia (Yirmiya, 1996; Bhattacharya et al., 2018; Scheggi et al., 2018). Behavioral phenotypes in these models typically present in two phases. The initial phase dominated by sickness behavior is followed by a second phase in which sickness phenotypes wane, but depressive-like behaviors remain (Dantzer et al., 2008). Similarly, we describe how NIM mice showed reduced motility on the first day and anhedonia measured as a lack of preference for sucrose for up to two days. It has been recently discovered, that a single subanesthetic dose of ketamine has an immediate and long lasting antidepressant effect in patients that are resistant to other treatments (Berman et al., 2000; Price et al., 2009). Interestingly, a single dose of ketamine was able to rescue both anhedonia and motor dysfunction in NIM mice, however fluoxetine had no effects in any of the measures we assessed. Increased inflammation often correlates with treatment resistance. Ketamine is an *N*-methyl-D-aspartate receptor (NMDAR) antagonist with well-documented actions through neuronal mechanisms (Kavalali, 2015). However, it is still not clear if ketamine has direct actions related to neuroinflammatory signaling, but there is emerging evidence for effects on astrocytes (Stenovec et al., 2020a; Stenovec et al., 2020b). Our use of ketamine is not contingent on knowing its precise mechanism, but does provide assurance that the anhedonia we tracked has relevance to depression-related phenotypes.

To better understand how neuroinflammation affected astrocytes during the sickness (Day 1), depressive-like (Day 2), and recovery (Day 14) phases we performed astrocyte RNA-seq from two cortical regions: the PFC, which is relevant to anhedonia and cognitive decline in many CNS diseases, and VCX, an area engaged in different visual circuits and behaviors. In parallel to the severity of the NIM phenotypes, the highest number of DEGs in both PFC and VCX were detected at Day 1, followed by Day 2. No DEGs were identified at Day 14. Transcriptionally different astrocyte populations have been observed between and within brain regions (Haim and Rowitch, 2017; Khakh and Deneen, 2019). We compared PFC and VCX astrocyte transcriptomes to assess if the response of astrocytes form two different brain circuits to the same stimulus was equivalent or region dependent. As expected, the transcriptomes of both PFC and VCX astrocytes changed during neuroinflammation. Interestingly, around 60% of the region-specific genes were common between control and NIM, indicating that a considerable proportion of identity defining gene expression in these two areas were preserved during neuroinflammation. When we focused on DEGs between the NIM vs control in each brain region, we discovered that one third were region specific. Our data show that astrocytes from distinct regions respond differently to the same stimulus, which merits further mechanistic work. This is potentially an important question with relevance to regional susceptibility of CNS regions to specific diseases. Furthermore, this result underscores the need for considerable caution when analyzing gene expression data that are available in various resources and published studies: the impact of regional variation (Haim and Rowitch, 2017; Khakh and Deneen, 2019), context-specific responses (Yu et al., 2020), and of variable responses to the same stimulus that is dependent on the brain region, as shown here, need to be thoughtfully accounted for in order to make meaningful conclusions. To what extent these region specific responses are due to cell autonomous mechanisms or to the variability of response of surrounding cells needs further exploration.

We evaluated cell-cell interaction mechanisms during neuroinflammation. To this end, we identified the brain cell-type enriched genes of the frontal and posterior cortices by using the datasets that Saunders and colleagues have generated (Saunders et al., 2018). We then asked which one of those genes changed their expression in our whole tissue (INPUT) samples in response to neuroinflammation. The cell type with the most upregulated genes was microglia followed by endothelial cells. Both microglia and endothelial cells also displayed clear activation of immune signaling. Interestingly, our cell-cell interaction analysis identified several ligand-receptor pairs that may signal from endothelial cells to microglia and astrocytes. Since endothelial cells are the first barrier between the blood and the brain at the neurovascular unit, our data suggest that these cells may translate peripheral inflammatory signals into the brain by directly signaling to astrocytes and microglia, potentially via the mechanisms that our data and analyses reveal.

It has been proposed that LPS induced neuroinflammation leads to release of C1q, TNF, and Il1α by microglia which in turn convert astrocytes into neurotoxic A1 astrocytes, which could lead to neuronal death in mouse models and human neurodegenerative disease (Liddelow et al., 2017). In this manuscript we used the same LPS dose and i.p. injection that was originally used to define a set of genes that characterize this reactive A1 phenotype (Zamanian et al., 2012). Accordingly, most of the panreactive and A1 genes were upregulated 1 day after LPS injection in both PFC and VCX. However, no neuronal death or neuronal electrophysiological dysfunction was observed in the PFC. Recently, it has been suggested that for reactive astrocytes to be neurotoxic, injury must precede the astrocyte transformation to A1 states for the neurons to be affected in the retina (Guttenplan et al., 2020). Additional studies are needed to better understand the nuances of any protective and detrimental roles of reactive astrocytes in pathology. Our studies suggest caution in concluding that neurons would be dysfunctional based on the appearance of published proposed correlative neurotoxic gene expression profiles within astrocytes. Our data also provide new avenues for exploration, such as studying how downregulation of cholesterol synthesis and loss of neuron support affects neural circuits (Boisvert et al., 2018; Itoh et al., 2018; Diaz-Castro et al., 2019; Mathys et al., 2019). The findings and gene expression data provide a rich source of information to formulate and test specific hypotheses in relation to the influence of neuroinflammation on brain function.

## Acknowledgments

Thanks to the UCLA Neuroscience Genomics Core for assistance with sequencing, and Fuying Gao for helping with RNA-seq data analysis. This work was supported by the National Institutes of Health (R35NS111583) and by the Ressler Family Foundation (BSK). B-DC was supported by Khakh lab funds and the UCLA Depression Grand Challenge. Collaborations between BSK and MVS groups were supported by the UCLA Depression Grand Challenge. AMB was supported by Sofroniew lab funds and the UCLA Depression Grand Challenge. We acknowledge the NINDS Informatics Center for Neurogenetics and Neurogenomics (P30 NS062691 to GC) and the Genetics, Genomics and Informatics Core of the Semel Institute of Neuroscience at UCLA (U54HD087101-01 from the Eunice Kennedy Shriver National Institute of Child Health and Human Development).

## Extended data

Extended data Figures 1-10

**Extended data Figure 1:**
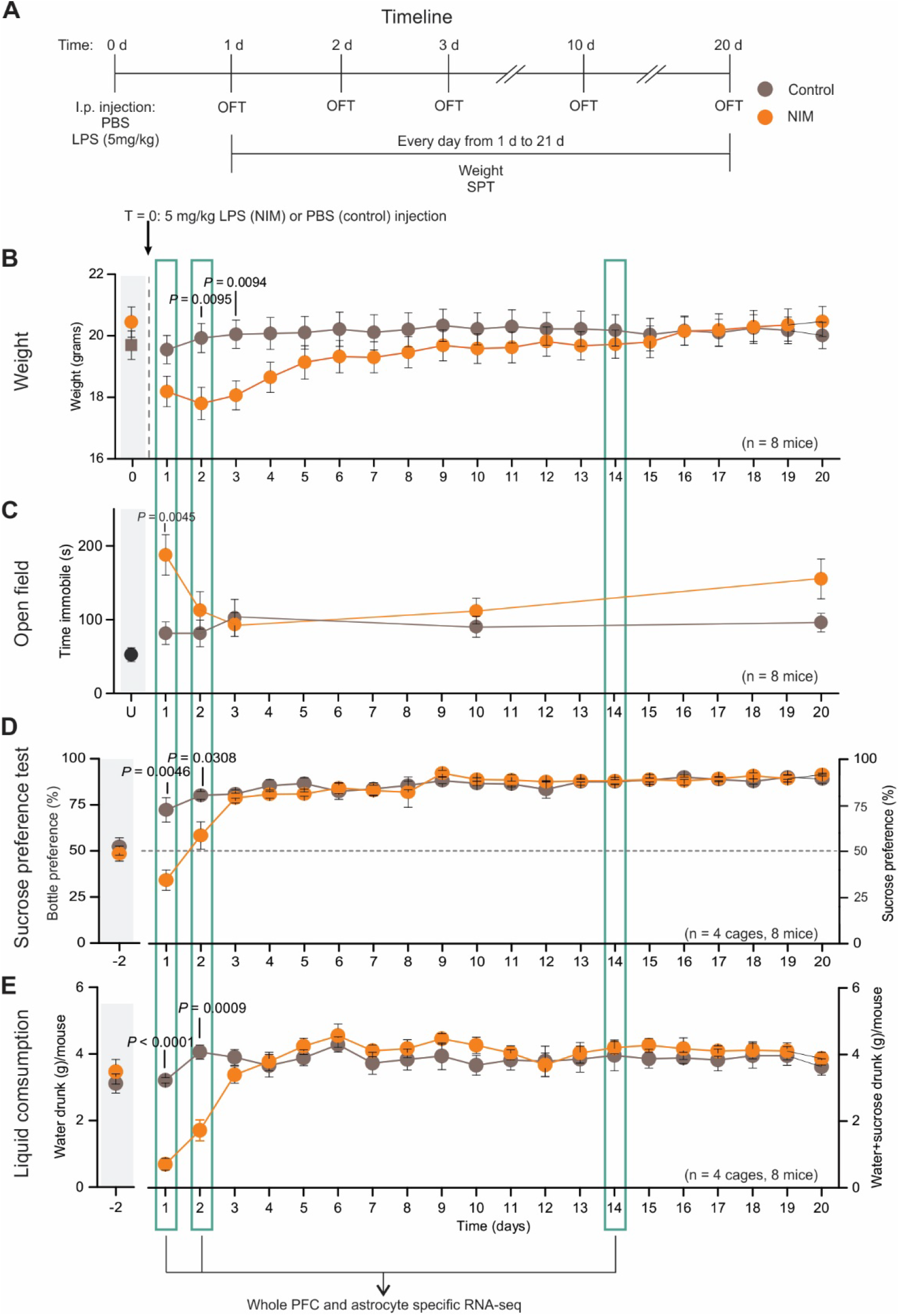
Behavioral assessment of LPS injected mice. ***A,*** Experimental design for LPS treatment and behavior assessment. ****B,**** Weight of control and LPS injected animals before injection (baseline) and during 20 d after injection. ****C,**** Time immobile during 6 minutes of open field test 1 d, 2 d, 3 d, 10 d and 20 d after injection. Baseline was generated with non-injected animals (“U”). ****D,**** On the left of the graph, the bottle preference when the mice are given the choice between two bottles of water two days before the injection (baseline). On the right, sucrose preference test during 20 d after injection. ****E,**** Total amount of liquid drunk per mouse two days before injection (baseline) and during 20 d after injection. Grey shading on the left, indicates baseline conditions from non-injected mice. Closed circles indicate mean ± s.e.m. In some cases, the error bars representing s.e.m are smaller than the symbol used for the mean.

**Extended data Figure 2:**
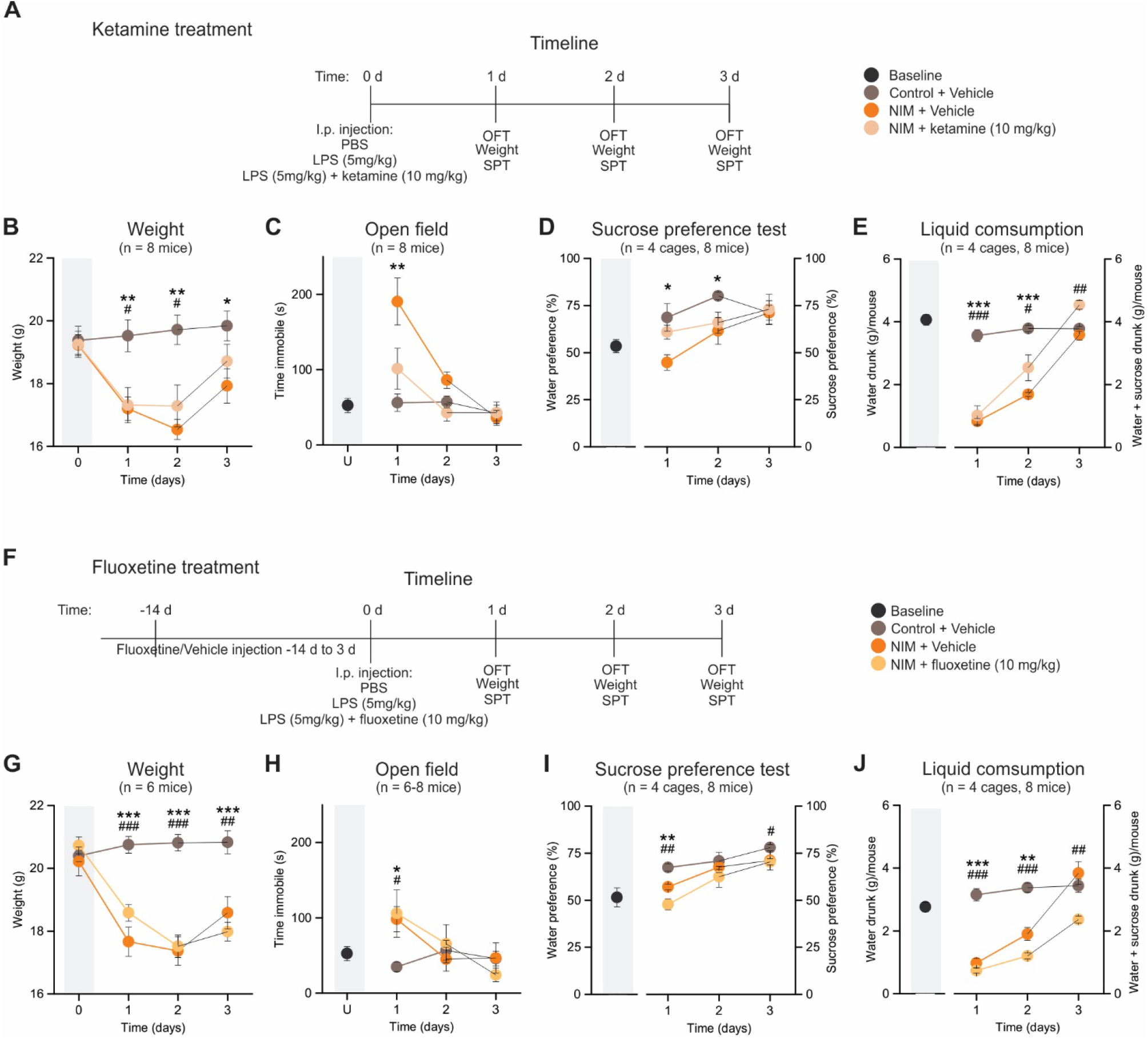
Behavioral assessment of LPS injected mice with reversal by ketamine and non-reversal by fluoxetine. ***A,*** Experimental design of LPS + ketamine treatment and behavior assessment. ***B,*** Weight of control, LPS and LPS + ketamine injected animals, before injection (baseline) and 1 d, 2 d and 3 d after injection. ***C,*** Time immobile during 6 minutes of open field test 1 d, 2 d and 3 d after injection. Baseline was generated with non-injected animals. ***D,*** On the left of the graph, the bottle preference when the mice are given the choice between two bottles of water two days before the injection (baseline). On the right, sucrose preference test 1 d, 2 d and 3 d after injection. ***E,*** Total amount of liquid drunk per mouse two days before injection (baseline), and 1 d, 2 d and 3 d after injection. ***F,*** Experimental design for LPS + fluoxetine treatment and behavior assessment. ***G,*** Weight of control, LPS and LPS + fluoxetine injected animals, before injection (baseline) and 1 d, 2 d and 3 d after injection. ***H,*** Time immobile during 6 minutes of open field test 1 d, 2 d and 3 d after injection. Baseline was generated with non-injected animals. ***I,*** On the left of the graph, the bottle preference when the mice are given the choice between two bottles of water two days before the injection (baseline). On the right, sucrose preference test 1 d, 2 d and 3 d after injection. ***J,*** Total amount of liquid drunk per mouse two days before injection (baseline), and 1 d, 2 d and 3 d after injection. Closed circles indicate mean ± s.e.m. In some cases, the error bars representing s.e.m are smaller than the symbol used for the mean. * indicates statistically significant difference between control and NIM and # between control and NIM + drug. * or ^#^ *P* < 0.05 > 0.01, ** or ^##^ *P* < 0.01 > 0.001, *** or ^###^ *P* < 0.001.

**Extended data Figure 3:**
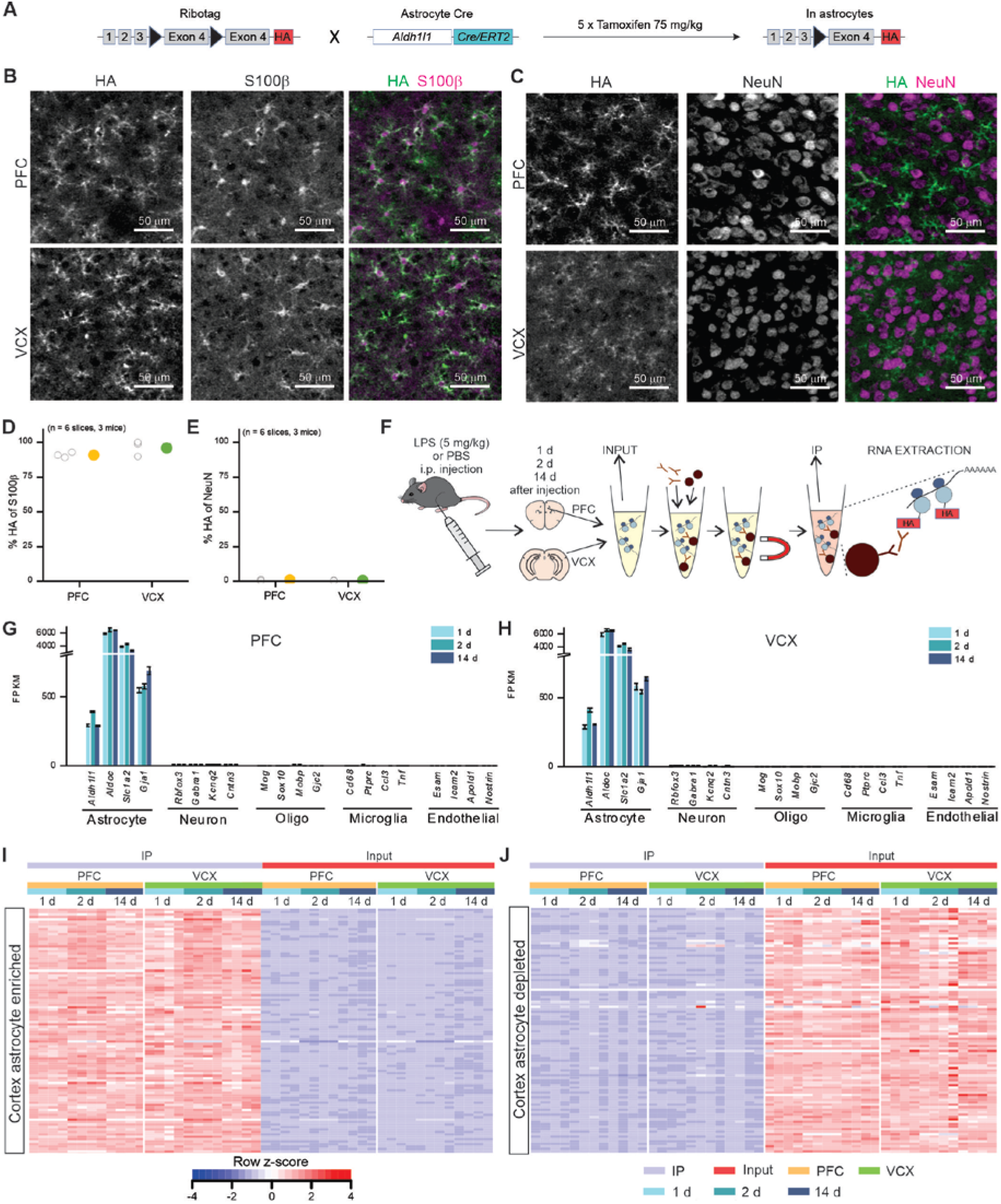
Prefrontal cortex (PFC) and visual cortex (VCX) whole tissue and astrocyte specific RNA purification for RNA-seq. ***A,*** Breeding strategy to obtain astrocyte specific expression of Rpl22HA by crossing Aldh1l1-Cre/ERT2 (Srinivasan et al. 2016) with Ribotag mice (Sanz et al. 2009). Cre excision of the Ribotag exon 4 is induced by injecting tamoxifen i.p. *B,* Representative images from HA and S100β staining in PFC and VCX. *C,* Representative images from HA and NeuN staining in PFC and VCX. *D,* % of S100β^+^ cells that express HA in PFC and VCX. *E,* % of NeuN^+^ cells that express HA in PFC and VCX. ****F,**** Sample collection method. PFC and VCX were dissected from LPS or PBS injected mice, one, two or 14 days after the injection. Tissue was homogenized, HA-tagged ribosomes immunoprecipitated, and RNA extracted from IP and input samples. ****G-H,**** RNA-seq gene expression level (FPKM) of four widely used markers for brain cell types in PFC (****G****) and VCX control samples (****H****). ****I-J,**** Heat map showing relative RNA-seq expression (z-score) of the top 100 adult-cortex astrocyte-enriched genes (enriched in IP vs input with FPKM > 10 (Srinivasan et al. 2016)) (****I****) and in PFC and top 100 adult whole-cortex enriched genes (depleted in IP vs input (Srinivasan et al. 2016)) (****J****) in PFC and VCX control samples. In the scatter plots, open circles are raw data with closed circles indicating mean ± s.e.m. In some cases, the error bars representing s.e.m are smaller than the symbol used for the mean.

**Extended data Figure 4:**
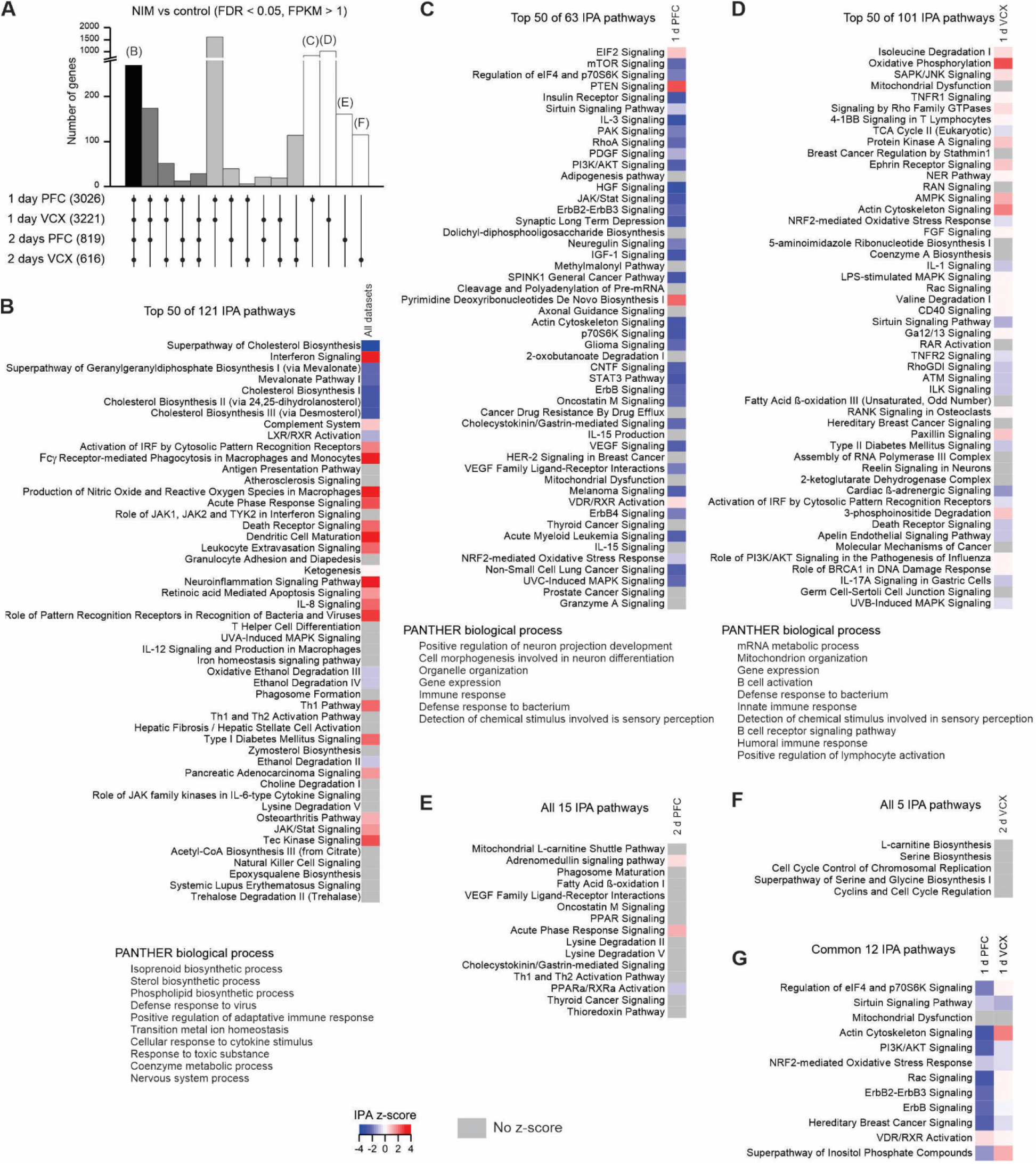
Pathways altered in NIM astrocytes. ***A,*** Overlap of astrocyte NIM vs control DEGs (FDR < 0.05, FPKM > 1) between PFC and VCX at 1 d and 2 d (from Figure 2M). ****B-D,**** Top 50 IPA canonical pathways (top) and PANTHER biological processes (bottom) identified for the NIM vs control DEGs common across 1 d PFC, 1 d VCX, 2 d PFC and 2 d VCX (****B****), unique for 1 d PFC (****C****) or unique for 1 d VCX (****D****). ****E-F,**** All IPA canonical pathways identified for the NIM vs control DEGs unique for 2 d PFC (****E****) or unique for 2 d VCX (****F****). ****G,**** Common IPA canonical pathways for the NIM vs control DEGs unique for 1 d PFC and 1 d VCX. IPA z-score indicates if the pathway is predicted to be inhibited (blue) or activated (red). In some cases, activation or inhibition cannot be predicted (grey).

**Extended data Figure 5:**
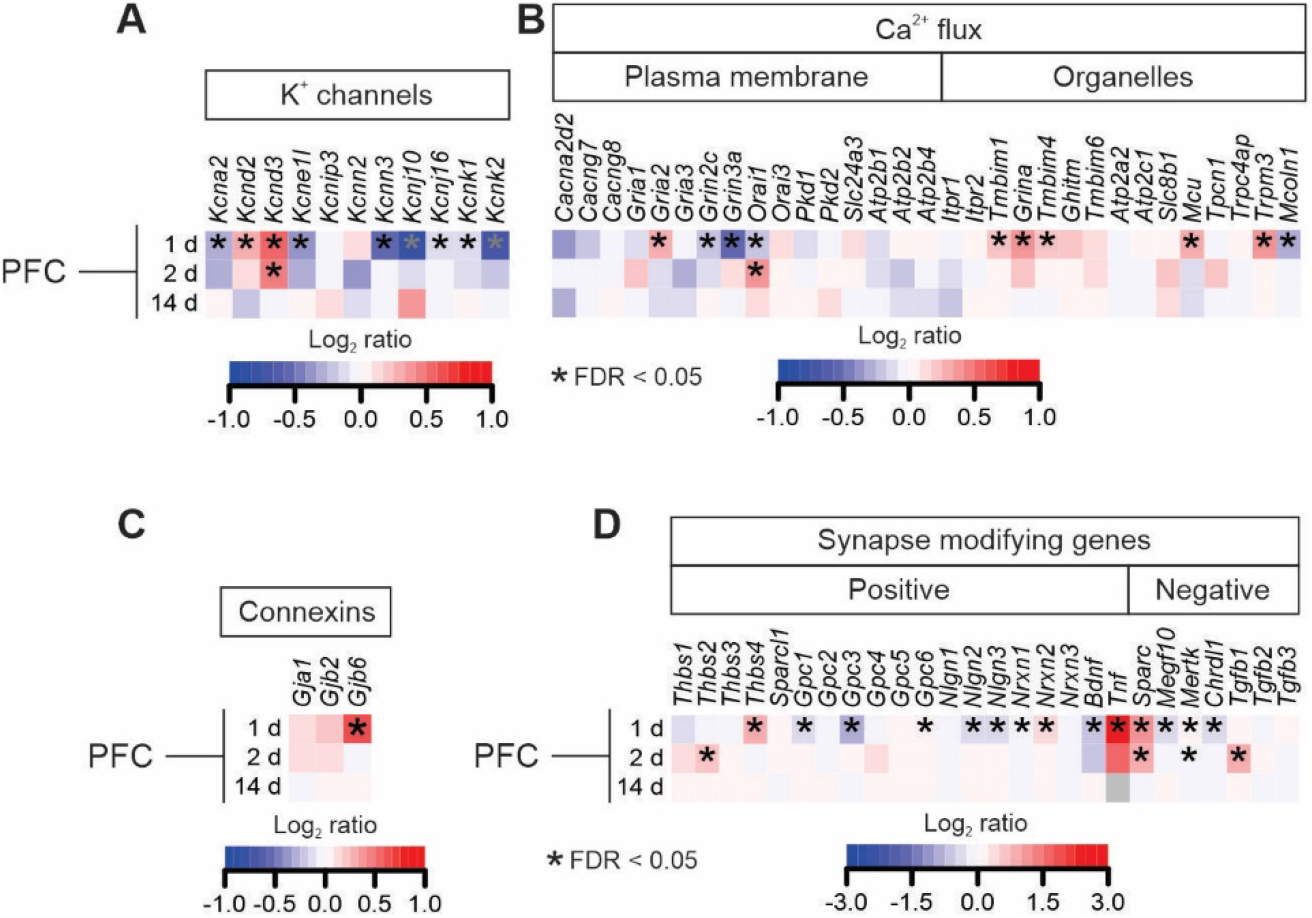
Alterations of the expression of genes involved in reactivity, K+ channels, Ca^2+^ flux, gap junctions and synapse modification in NIM PFC. *A,* NIM vs control differential expression of the genes encoding K^+^ channels (with an FPKM > 10) in 1 d, 2 d and 14 d PFC astrocytes. *B,* NIM vs control differential expression of the genes encoding proteins involved in Ca^2+^ flux (with an FPKM > 10) in 1 d, 2 d and 14 d PFC astrocytes. *C,* NIM vs control differential expression of the genes encoding connexins (with an FPKM > 10) in 1 d, 2 d and 14 d PFC astrocytes. *D,* NIM vs control differential expression of the genes encoding proteins involved synapse modification in 1 d, 2 d and 14 d PFC astrocytes. In the heat maps, red means upregulation and blue downregulation when comparing NIM vs control. * indicates the genes that were differentially expressed with FDR < 0.05.

**Extended data Figure 6:**
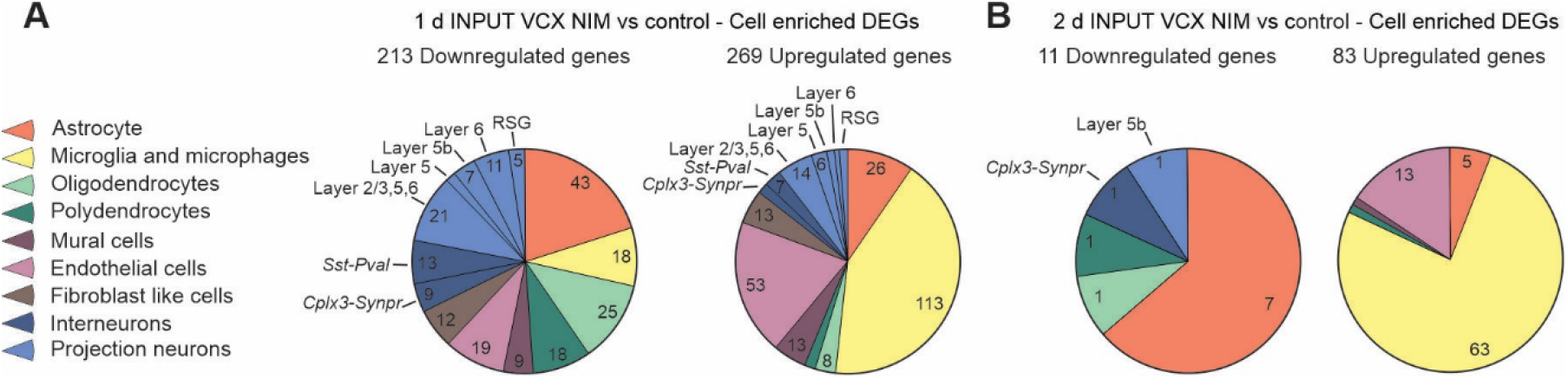
VCX cell specific DEG contribution using input NIM vs control RNA-seq. ***A,*** Pie charts indicating the proportion of 1 d input downregulated (left) and upregulated (right) genes that are specific for each cell type listed in the legend on the left. The numbers in each section of the pie chart indicate the number of DEGs for that cell type. ****B,**** Same as in A but for 2 d input samples.

**Extended data Figure 7:**
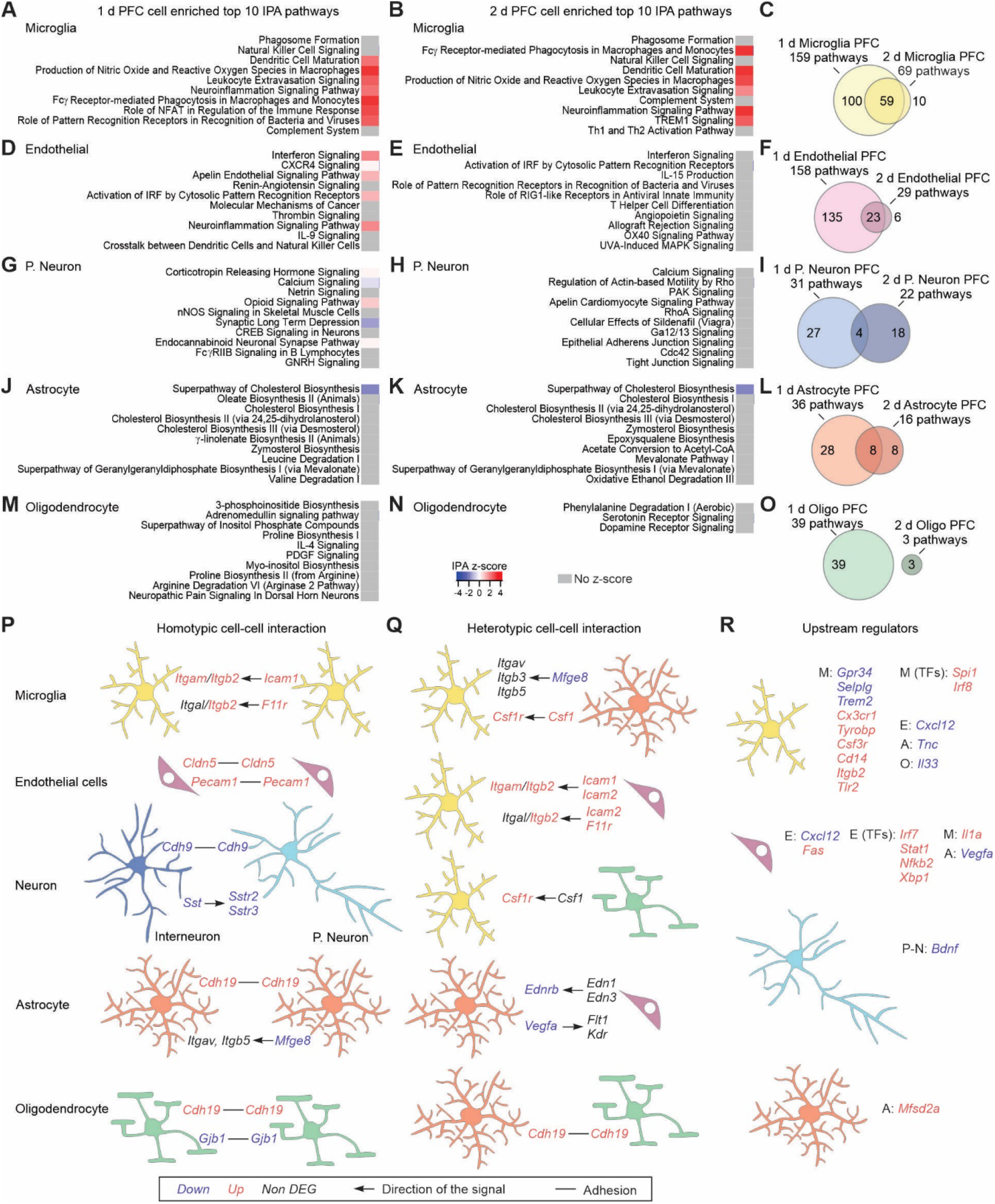
PFC cell specific altered pathways and cell-cell interaction mechanisms. ***A-B,*** Top 10 IPA pathways identified for the microglia enriched DEGs at 1 d (A) or 2 d (*B*). *C,* Overlap of the identified IPA pathways from microglia enriched DEGs between 1 d and 2 d. ****D-O,**** Same as ****A-C**** but for endothelial cells (****D-F****), projection neurons (****G-I****), astrocytes (****H-L****), or oligodendrocytes (****M-O****). ****P-Q,**** Homotypic (****P****) and heterotypic (****Q****) adhesion and ligand-receptor cell-cell interaction mechanisms that are altered in NIM. The arrows indicate the directionality of the signal, e.g. ligand → receptor. ****R,**** Differentially expressed upstream regulators of microglia, endothelial, projection neuron and astrocyte DEGs. The upstream regulators could be receptors, ligands, or transcription factors (TFs) that are expressed in the same cell, or in the neighboring ones: microglia (M), endothelial (E), projection neuron (P-N), astrocyte (A) or oligodendrocyte (O).

**Extended data Figure 8:**
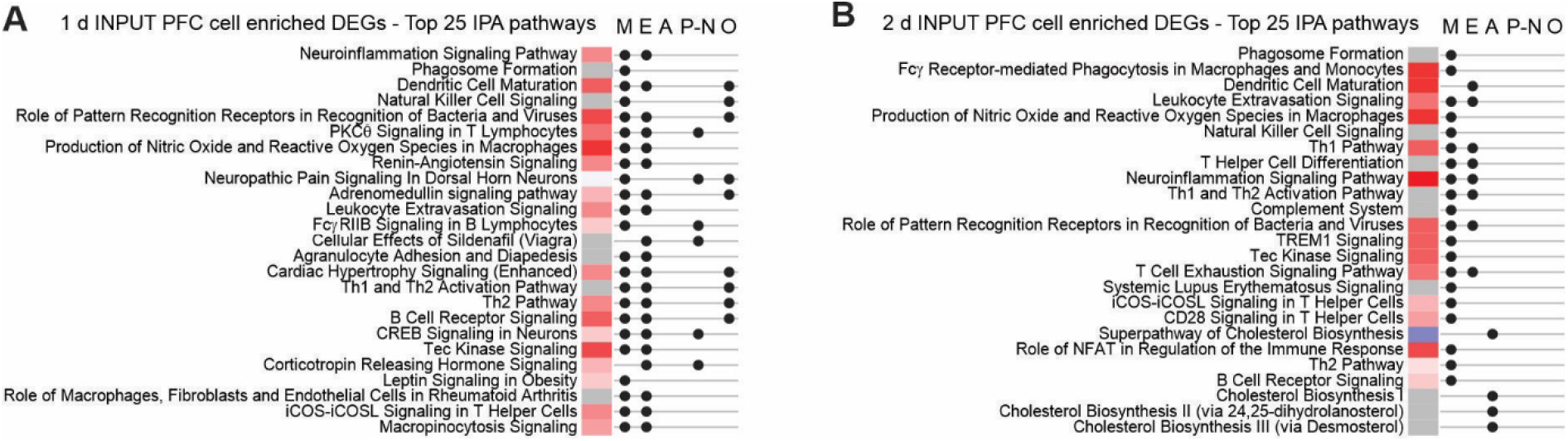
PFC altered pathways in specific cell types. ***A-B,*** Top 25 Ingenuity Pathway Analysis (IPA) pathways identified for the cell specific DEGs at 1 d (****A****) or 2 d (****B****). The heat map indicates in red the pathways that are activated and in blue the ones that are inhibited. On the right of the heat maps the dots indicate the pathways that were also found when performing IPA only with microglia (M), endothelial (E), projection neuron (P-N), astrocyte (A) and oligodendrocyte (O) specific DEGs.

**Extended data Figure S9:**
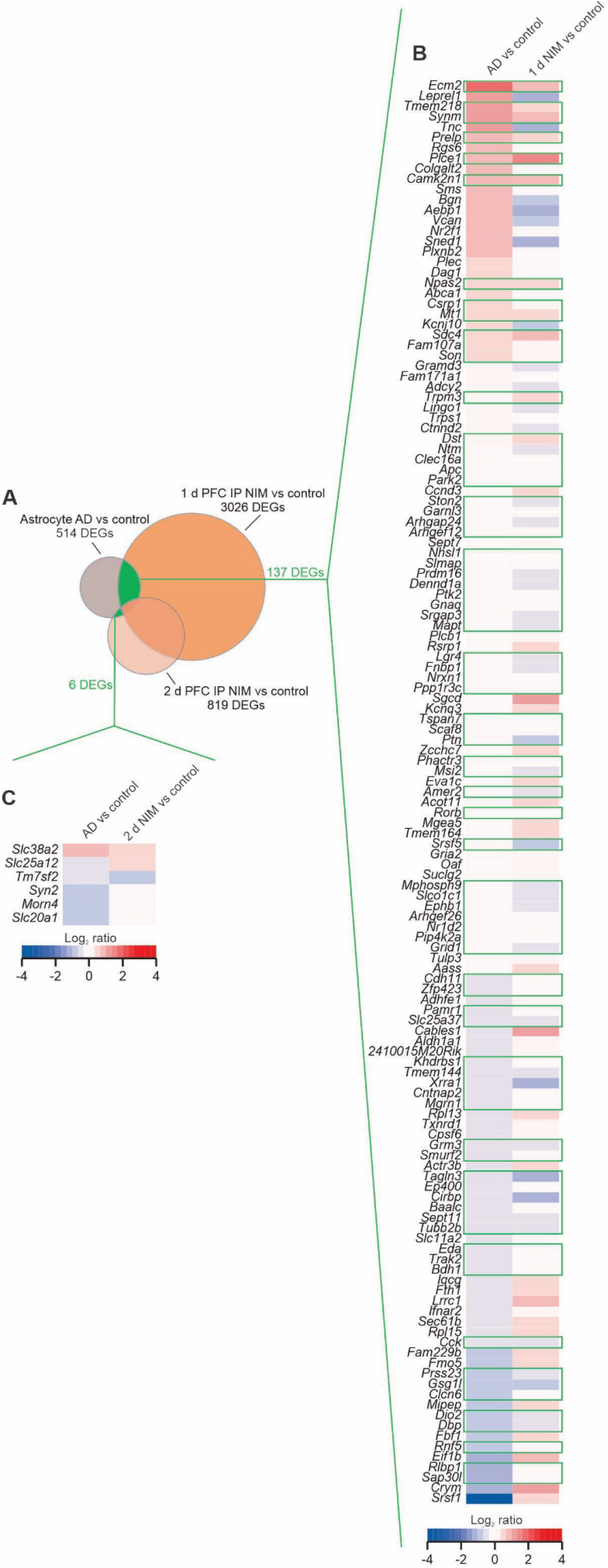
Comparison of PFC transcriptional profiles between 1 d and 2 d NIM and human AD. ***A,*** Prefrontal cortex snRNA-seq data from 24 AD patients and 24 controls identified 514 astrocyte DEGs (Mathys et al., 2019). ***B,*** Heatmap showing the differential expression magnitude in log_2_ ratio of the 137 common astrocyte DEGs between AD and 1 d NIM but not differentially expressed at 2 d NIM. ***C,*** Heatmap showing the differential expression magnitude in log_2_ ratio of the 6 common astrocyte DEGs between AD and 2 d NIM but not differentially expressed at 1 d NIM. In the heat maps, red means upregulation and blue downregulation when comparing the condition of study vs its control. The green frames on the heatmaps highlight the genes that change in the same direction in the human disease and mouse neuroinflammation.

**Extended data Figure 9b:**
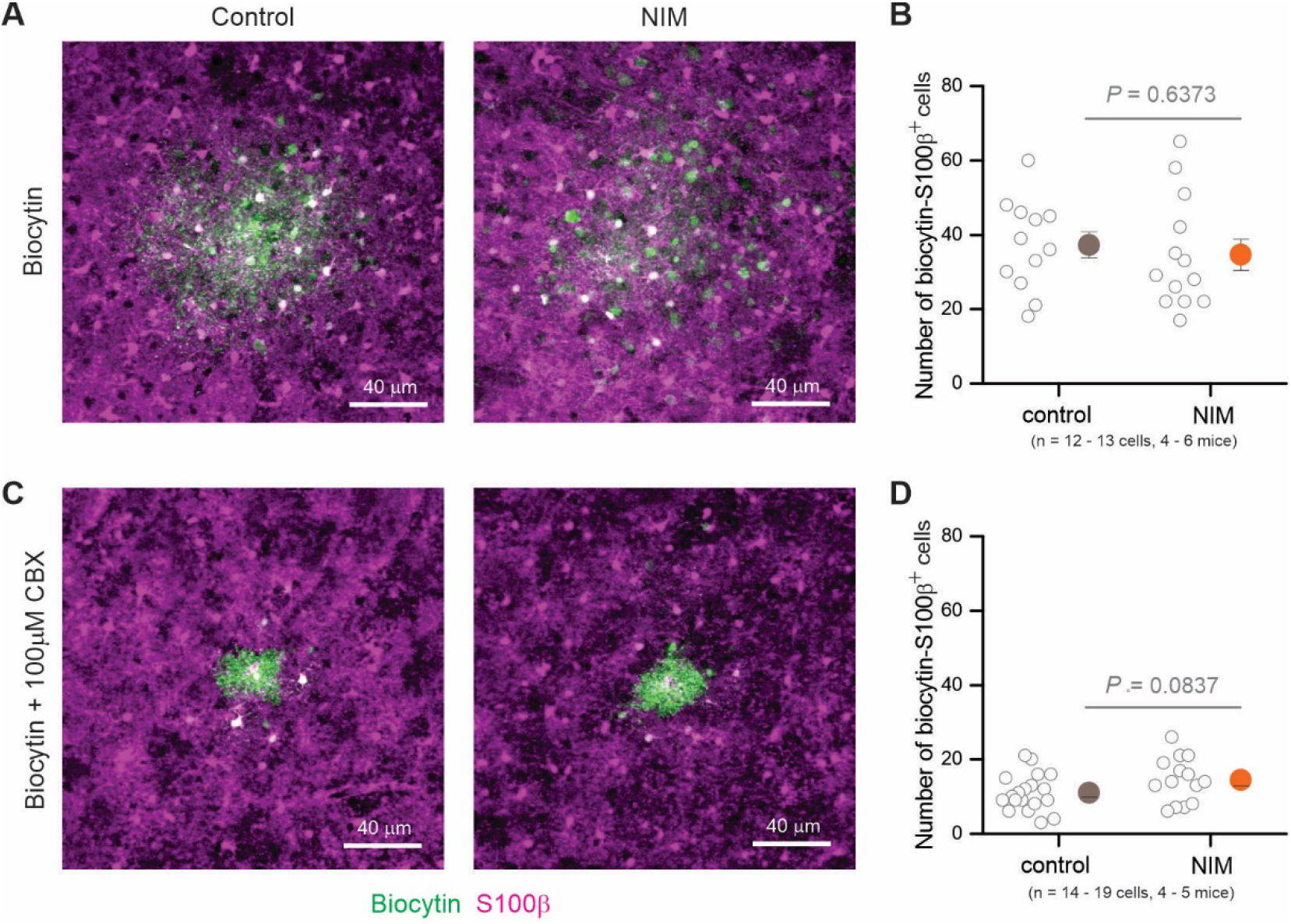
Cell coupling measured as diffusion of biocytin between astrocytes. ***A,*** Representative images from biocytin and S100β staining in PFC astrocytes filled in aCSF. ****B,**** Number of S100β^+^ cells that were filled with biocytin. (C) Representative images from biocytin and S100β staining in PFC astrocytes filled in aCSF + 100 μM CBX. (D) Number of S100β^+^ cells that were filled with biocytin in CBX. Open circles are raw data with closed circles indicating mean ± s.e.m. In some cases, the error bars representing s.e.m are smaller than the symbol used for the mean.

